# Tuning site-specific dynamics to drive allosteric activation in a pneumococcal zinc uptake regulator

**DOI:** 10.1101/301853

**Authors:** Daiana A. Capdevila, Fidel Huerta, Katherine A. Edmonds, My T. Le, Hongwei Wu, David P. Giedroc

**Affiliations:** Department of Chemistry, Indiana University, 800 E. Kirkwood Drive, Bloomington, IN 47405-7102, United States; Graduate Program in Biochemistry, Indiana University, 212 S. Hawthorne Drive, Bloomington, IN 47405, United States; Department of Molecular and Cellular Biochemistry, Indiana University, 212 S. Hawthorne Drive, Bloomington, IN 47405 United States

**Author notes:** Correspondence to David P. Giedroc.

## Abstract

MarR (multiple antibiotic resistance repressor) family proteins are bacterial repressors that regulate transcription in response to a wide range of chemical signals. Although specific features of MarR family function have been described, the role of atomic motions in MarRs remains unexplored thus limiting insights into the evolution of allostery in this ubiquitous family of repressors. Here, we provide the first experimental evidence that internal dynamics play a crucial functional role in MarR proteins. Streptococcus pneumoniae AdcR (adhesin-competence repressor) regulates Zn^II^ homeostasis and Zn^II^ functions as an allosteric activator of DNA binding. Zn^II^ coordination triggers a transition from independent domains to a more compact structure. We identify residues that impact allosteric activation on the basis of Zn^II^-induced perturbations of atomic motions over a wide range of timescales. These findings reconcile the distinct allosteric mechanisms proposed for other MarRs and highlight the importance of conformational dynamics in biological regulation.

## Introduction

Successful bacterial pathogens respond to diverse environmental insults or changes in intracellular metabolism by modulating gene expression (Alekshun & Levy, 2007). Such changes in gene expression are often mediated by “one-component” transcriptional regulators, which directly sense chemical signals and convert such signals into changes in transcription. Members of the multiple antibiotic resistance regulator (MarR) family are critical for the survival of pathogenic bacteria in hostile environments, particularly for highly antibiotic-resistant pathogens (Ellison & Miller, 2006, Yoon *et al.*, 2009, Weatherspoon-Griffin & Wing, 2016, Tamber & Cheung, 2009, Aranda *et al.*, 2009, Grove, 2017). Chemical signals sensed by MarRs include small molecule metabolites (Deochand & Grove, 2017), reactive oxygen species (ROS) (Liu *et al.*, 2017, Sun *et al.*, 2012) and possibly reactive sulfur species (RSS) (Peng *et al.*, 2017). It has been proposed that evolution of new MarR proteins enables micro-organisms to colonize new niches (Deochand & Grove, 2017), since species characterized by large genomes and a complex lifestyle encode many, and obligate parasitic species with reduced genome sizes encode few (Perez-Rueda *et al.*, 2004). Therefore, elucidating how new inducer specificities and responses have evolved in this ubiquitous family of proteins on what is essentially an unchanging molecule scaffold is of great interest, as is the molecular mechanism by which inducer binding or cysteine thiol modification allosterically regulates DNA operator binding in promoter regions of regulated genes.

Obtaining an understanding of how allostery has evolved in one-component regulatory systems (Ulrich *et al.*, 2005, Marijuan *et al.*, 2010), including MarR family repressors, requires a comprehensive analysis of the structural and dynamical changes that occur upon inducer and DNA binding (Capdevila *et al.*, 2017a, Tzeng & Kalodimos, 2013, West *et al.*, 2012, Tzeng & Kalodimos, 2009). For MarRs, several distinct allosteric mechanisms have been proposed, from a “domino-like” response (Bordelon *et al.*, 2006, Gupta & Grove, 2014, Perera & Grove, 2010) to ligand binding-mediated effects on asymmetry within the dimer (Anandapadamanaban *et al.*, 2016), to oxidative crosslinking of *E. coli* MarR dimers into DNA binding-incompetent tetramers (Hao *et al.*, 2014). While there are more than 130 crystal structures of MarR family repressors in different allosteric states (Fig. S1), an understanding of the role of atomic motions and the conformational ensemble in MarRs is nearly totally lacking and what is known is based exclusively on simulations (Anandapadamanaban *et al.*, 2016, Sun *et al.*, 2012). Here, we provide the first experimental evidence in solution that internal dynamics play a crucial functional role in a MarR protein, thus define characteristics that may have impacted the evolution of new biological outputs in this functionally diverse family of regulators.

In the conventional regulatory paradigm, the binding of a small molecule ligand, or the oxidation of conserved ROS-sensing cysteines induces a structural change in the homodimer that typically negatively impacts DNA binding affinity. This results in a weakening or dissociation of the protein-DNA complex and transcriptional derepression. Several reports provide evidence for a rigid body reorientation of the two α4 (or αR)-reading heads within the dimer (Fig. 1A-B, Fig. S1) (Alekshun *et al.*, 2001, Fuangthong & Helmann, 2002, Wilke *et al.*, 2008, Chang *et al.*, 2010, Liu *et al.*, 2017, Deochand & Grove, 2017, Dolan *et al.*, 2011, Deochand *et al.*, 2016).

**Figure 1.**
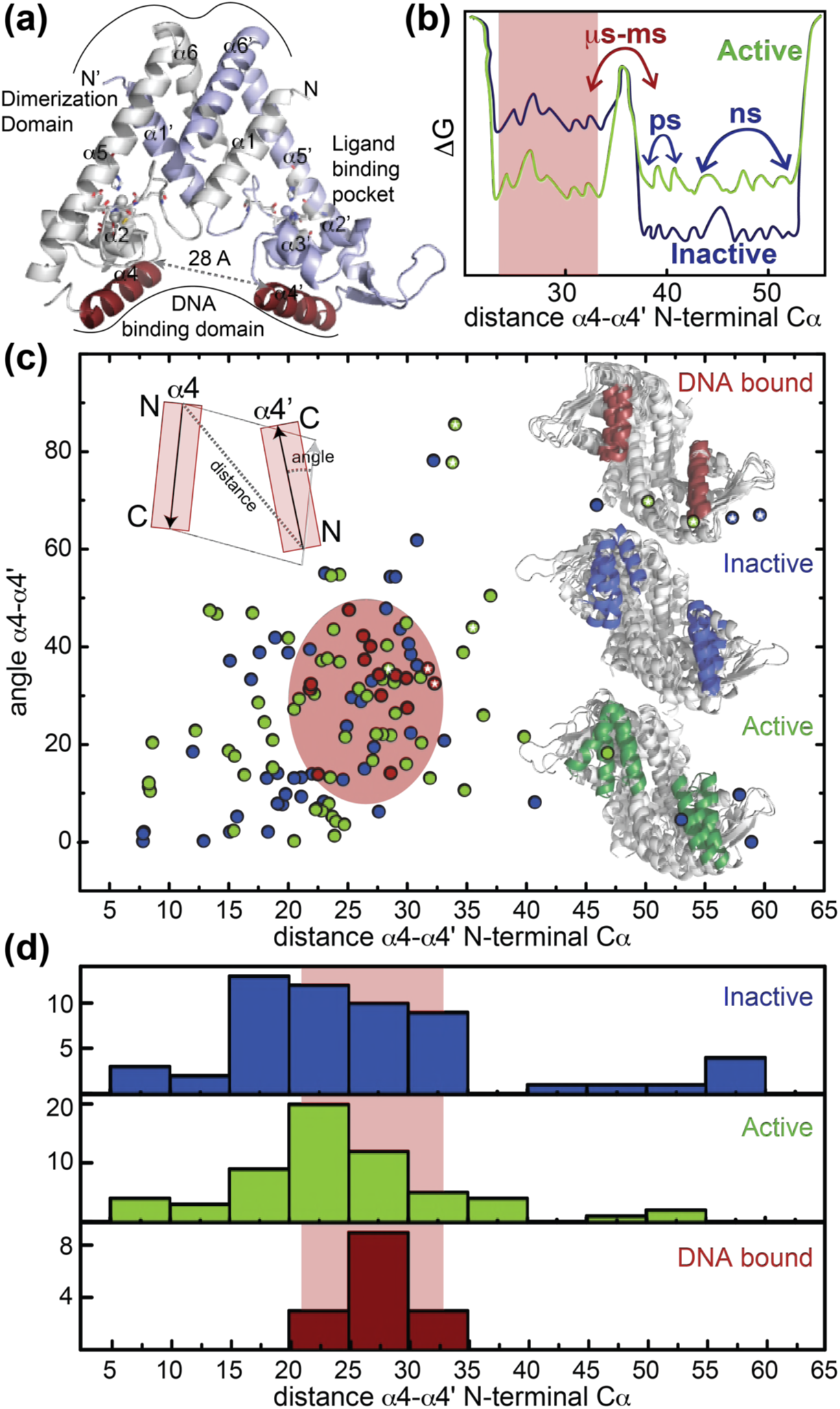
(a) Ribbon representation of dimeric Zn(II)-bound AdcR, with one protomer shaded white and the other shaded *light blue* (PDB-ID: 3tgn) (Guerra *et al.*, 2011). The two Zn(II) ions in each protomer are represented by spheres, and coordinating ligands are shown in stick representation. The DNA binding helices are shaded *red.* (b) Simplified free energy diagram showing of the active (*green*) and inactive (*blue*) states the relative population of two distinct conformations: compatible with DNA binding (*red* rectangle, α4-α4’ distance between DNA binding helices, ≈30 Å) and incompatible with DNA binding (larger α4-α4’ distances). (c) The α4-α4’ distance distribution plotted against the DNA-binding inter-helical α4-α4’ orientation distribution for all the reported MarR crystal structures (see Table S1 for details) in the allosterically “active” DNA-binding conformation (*green*), an “inactive” conformation (*blue*) and in the DNA-bound (*red*) conformation. The inferred conformational space occupied by the DNA-bound conformation in all MarR regulators (Table S1) is shaded in *red* oval. Ribbon representation of the molecules in each conformation are shown in the inset, as well as a scheme of how the distances and angles were measured. (d) Histogram plot of the α4-α4’ distance (see panel c) extracted from 136 different crystal structures of MarR repressors in the DNA-binding- “inactive”, DNA-binding “active” and DNA-bound conformations.

The generality of this simple paradigm is inconsistent with the findings that some MarR proteins share very similar static structures in the active (DNA binding-competent) and inactive (DNA binding-incompetent) states (Anandapadamanaban *et al.*, 2016, Kim *et al.*, 2016, Liguori *et al.*, 2016); furthermore, several active states have been shown to require a significant rearrangement to bind DNA (Alekshun *et al.*, 2001, Liu *et al.*, 2017, Zhu *et al.*, 2017b, Hao *et al.*, 2013, Gao *et al.*, 2017, Chin *et al.*, 2006, Saridakis *et al.*, 2008). In fact, a comprehensive analysis of all available MarR family structures strongly suggests that the degree of structural reorganization required to bind DNA, characterized by a narrow distribution of α4-α4’ orientations, is comparable whether transitioning from the inactive *or* active states of the repressor (Fig. 1C, Table S1). These observations strongly implicate a conformational ensemble model of allostery (Motlagh *et al.*, 2014) (Fig. 1B-D), where inducer sensing impacts DNA binding by restricting the conformational spread of the active repressor, as was proposed in a recent molecular dynamics study (Anandapadamanaban *et al.*, 2016).

MarR proteins are obligate homodimers that share a winged-helical DNA-binding domain connected to a DNA-distal all-helical dimerization domain where organic molecules bind in a cleft between the two domains (Fig. S1B). Individual MarR members have been shown to bind a diverse range of ligands at different sites on the dimer (Otani *et al.*, 2016, Takano *et al.*, 2016); likewise, oxidation-sensing cysteine residues are also widely distributed in the dimer (Fuangthong & Helmann, 2002, Liu *et al.*, 2017, Hao *et al.*, 2014, Dolan *et al.*, 2011, Chen *et al.*, 2006). This functional diversity is accompanied by relatively low overall sequence similarity, which suggests that a conserved molecular pathway that connects sensing sites and the DNA binding heads is highly improbable. Complicating our current mechanistic understanding of this family is that for many members, including *E. coli* MarR, the physiological inducer (if any) is unknown, rendering functional conclusions on allostery from crystallographic experiments alone less certain (Hao *et al.*, 2014, Zhu *et al.*, 2017b).

In contrast to the extraordinary diversity of thiol-based switching MarRs, MarR family metallosensors are confined to a single known regulator of Zn^II^ uptake, exemplified by AdcR (adhesin competence regulator) from *S. pneumoniae* and closely related *Streptococcus ssp.* (Loo *et al.*, 2003, Reyes-Caballero *et al.*, 2010) and ZitR from *Lactococcus spp* (Llull *et al.*, 2011, Zhu *et al.*, 2017c). AdcR and ZitR both possess two closely spaced pseudotetrahedral Zn^II^ binding sites termed site 1 and site 2 (Fig. 1A) that bind Zn^II^ with different affinities (Reyes-Caballero *et al.*, 2010, Guerra *et al.*, 2011, Sanson *et al.*, 2015, Zhu *et al.*, 2017c). Zn^II^ is an allosteric *activator* of DNA operator binding which is primarily dependent on the structural integrity of site 1 (Reyes-Caballero *et al.*, 2010, Zhu *et al.*, 2017c). ZitR has been recently extensively structurally characterized, with crystallographic models now available for the apo- and Zn^II^_1_- (bound to site 1) and Zn^II^_2_- and Zn^II^_2_-DNA operator complexes, thus providing significant new insights into ZitR and AdcR function (Zhu *et al.*, 2017c). These structures reveal that Zn^II^_2_-ZitR and Zn^II^_2_-AdcR form triangularly-shaped homodimers and are essentially identical, as anticipated from their high sequence identity (49%). Apo-ZitR adopts a conformation that is incompatible with DNA binding, and filling of both Zn^II^ sites is required to adopt a conformation that is similar to that of the DNA-complex. Thermodynamically, filling of the low affinity site 2 enhances allosteric activation of DNA-binding by ≈10-fold, and this occurs concomitant with a change in the H42 donor atom to the site 1 Zn^II^ ion from Nε2 in the apo- and Zn^II^_1_-states to Nδ1 in the Zn^II^_2_-ZitR (as in Zn^II^_2_ AdcR; (Guerra *et al.*, 2011)) and Zn^II^_2_ ZitR-DNA operator complexes (Zhu *et al.*, 2017c). Allosteric *activation* by Zn^II^ is in strong contrast to all other members of the MarR superfamily, consistent with its biological function as uptake repressor at high intracellular Zn^II^

Here we employ a combination of NMR-based techniques and small angle x-ray scattering (SAXS) to show that apo- (metal-free) AdcR in solution is characterized by multiple independent domains connected by flexible linkers, resulting in a distinct quaternary structure from the Zn-bound state previously structurally characterized (Guerra *et al.*, 2011). Our backbone relaxation dispersion-based NMR experiments show that apo-AdcR samples distinct conformational states in the μs-ms timescale, while Zn^II^ narrows this distribution by conformational selection, increasing the population of a state that has higher affinity for DNA. This finding is fully consistent with the crystallographic structures of Zn^II^_2_ ZitR and the Zn^II^_2_ ZitR:DNA complex (Zhu *et al.*, 2017c). The site-specific backbone and methyl sidechain dynamics in the ps-ns timescale show that Zn^II^ not only induces a general restriction of these protein dynamics, but also enhances fast timescale, low-amplitude motions in the DNA binding domains. Together, these data reveal that Zn^II^ coordination promotes a conformational change that reduces the entropic cost of DNA binding and enhances internal dynamics uniquely within the DNA binding domain, thus poising the repressor to interact productively with various DNA operator target sequences (Reyes-Caballero *et al.*, 2010). We demonstrate the predictive value of this allosteric model by functionally characterizing “cavity” mutants of AdcR (Capdevila *et al.*, 2017a). Overall, our findings suggest that protein dynamics on a wide range of timescales strongly impact AdcR function. This ensemble model of allostery successfully reconciles the distinct mechanisms proposed for other MarR family repressors and suggests a mechanism of how evolution tunes dynamics to render distinct biological outputs (allosteric activation vs. allosteric inhibition) on a rigorously conserved molecular scaffold.

## Results and Discussion

### Solution structural differences between apo and Zn^II^ bound forms of AdcR

Our crystal structure suggests that once AdcR is bound to both Zn^II^, the αR- (α4) reading heads adopt a favorable orientation for DNA binding (Guerra *et al.*, 2011), a finding fully compatible with structural studies of *L. lactis* ZitR (Zhu *et al.*, 2017c) (Fig. 1A). These structural studies suggest a “pre-locked” model, where Zn^II^ binding to both sites 1 and 2, concomitant with a H42 ligand atom switch, locks the AdcR homodimer into a DNA binding-competent conformation. This model makes the prediction that the unligated AdcR can explore conformations structurally incompatible with DNA binding, as shown previously for Zn^II^_1_ ZitR (Zhu *et al.*, 2017c), thus requiring a significant degree of reorganization to bind with high affinity to the DNA (Fig. 1B). Despite significant efforts, it has not yet been possible to obtain the crystal structure of apo-AdcR, suggesting that the apo-repressor may be highly flexible in solution (Guerra *et al.*, 2011, Sanson *et al.*, 2015). Thus, we employed SAXS as a means to explore the apo-AdcR structure and elucidate the structural changes induced by Zn^II^ binding and conformational switching within the AdcR homodimer.

We first examined the behavior of apo- and Zn^II^-bound states. Both states show Guinier plots indicative of monodispersity and similar radii of gyration (*R*_g_). These data reveal that each state is readily distinguished from the other in the raw scattering profiles (to *q*=0.5 Å^−1^) as well as in the PDDF plots (*p(r) versus r*), with the experimental scattering curve of the Zn^II^ bound state being consistent with one calculated from the Zn^II^_2_ AdcR crystal structure (Fig. 2A). Moreover, a qualitative analysis of the PDDF plots suggests that apo-AdcR is less compact than the Zn^II^-bound state (Fig. S2). The molecular scattering envelopes calculated as bead models with the *ab initio* program DAMMIF for apo-AdcR suggest that the differences between the apo and Zn^II^ AdcR SAXS profiles can be explained on the basis of a reorientation of the winged helix-turn-helix motif with respect to the dimerization domain, particularly in a distortion in the α5 helix (Fig. 2B). In an effort to obtain higher resolution models, we reconstructed atomic models from perturbations in the Zn-bound crystal structure that better fit the complete SAXS profiles (*q*<1.0). The models obtained confirm that the Zn-bound structure in solution resembles the crystallographic models of apo-ZitR and Zn^II^ AdcR (Guerra *et al.*, 2011, Zhu *et al.*, 2017c); however, we note that the SAXS profile of the apo-AdcR differs significantly from the ZitR crystal structure (Fig. S2E) which is likely related to the high flexibility of this state in solution. Moreover, the resolution of SAXS based models cannot be used to obtain residue-specific information about structural perturbations introduced by Zn^II^ binding (Fig. S2). Thus, we turned to NMR-based techniques to provide both high resolution and site-specific information on this highly dynamic system.

**Figure 2.**
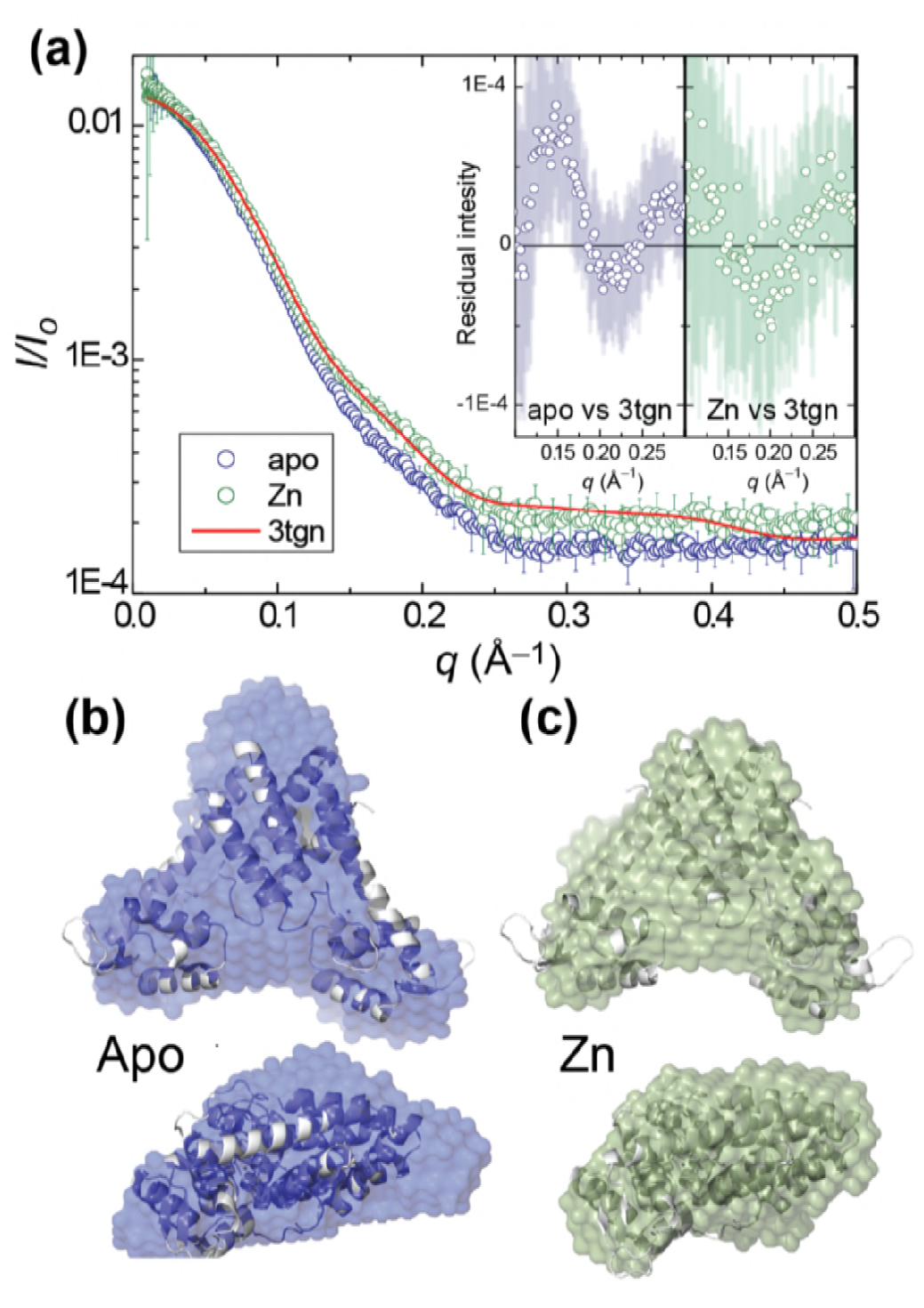
(a) Small angle X-ray scattering (SAXS) curve of AdcR in apo- and Zn_2_-states, with a curve calculated from the previously published AdcR-Zn_2_ structure (Fig. 1A, PDB-ID: 3tgn) (Guerra *et al.*, 2011). Best-fit DAMMIF *ab initio* model(Franke & Svergun, 2009) for apo- (b) (*blue*) and Zn^II^_2_-states (c) (*green*), aligned with the ribbon representation of the Zn^II^_2_ structure (Fig. 1A, PDB-ID: 3tgn). The corresponding Guinier, Kratky and pairwise distribution histogram plots are shown in Fig. S2, along with the fitting parameters.

TROSY NMR on 100% deuterated AdcR and optimized buffer conditions for both states (pH 5.5, 50 mM NaCl, 35 °C) enabled us to obtain complete backbone assignments for Zn^II^-AdcR and nearly complete for apo-AdcR (missing residues 21, 38-40 due to exchange broadening, Fig. 3). The chemical shift perturbation maps (Fig. 3A-B) reveal that the largest perturbations are found in the structural vicinity of the metal site region, *i.e*, the α1-α2 loop (residues 21-35), the remainder of the α2 helix (residues 41-47), and the central region of the α5 helix, which provides donor groups to both site 1 (H108, H112) and site 2 (E107) Zn^II^. These changes derive from changes in secondary structure, such as the extension of the α1 helix and partial unfolding of the α2 helix (Fig. S3), as well as from proximity to the Zn^II^.

**Figure 3.**
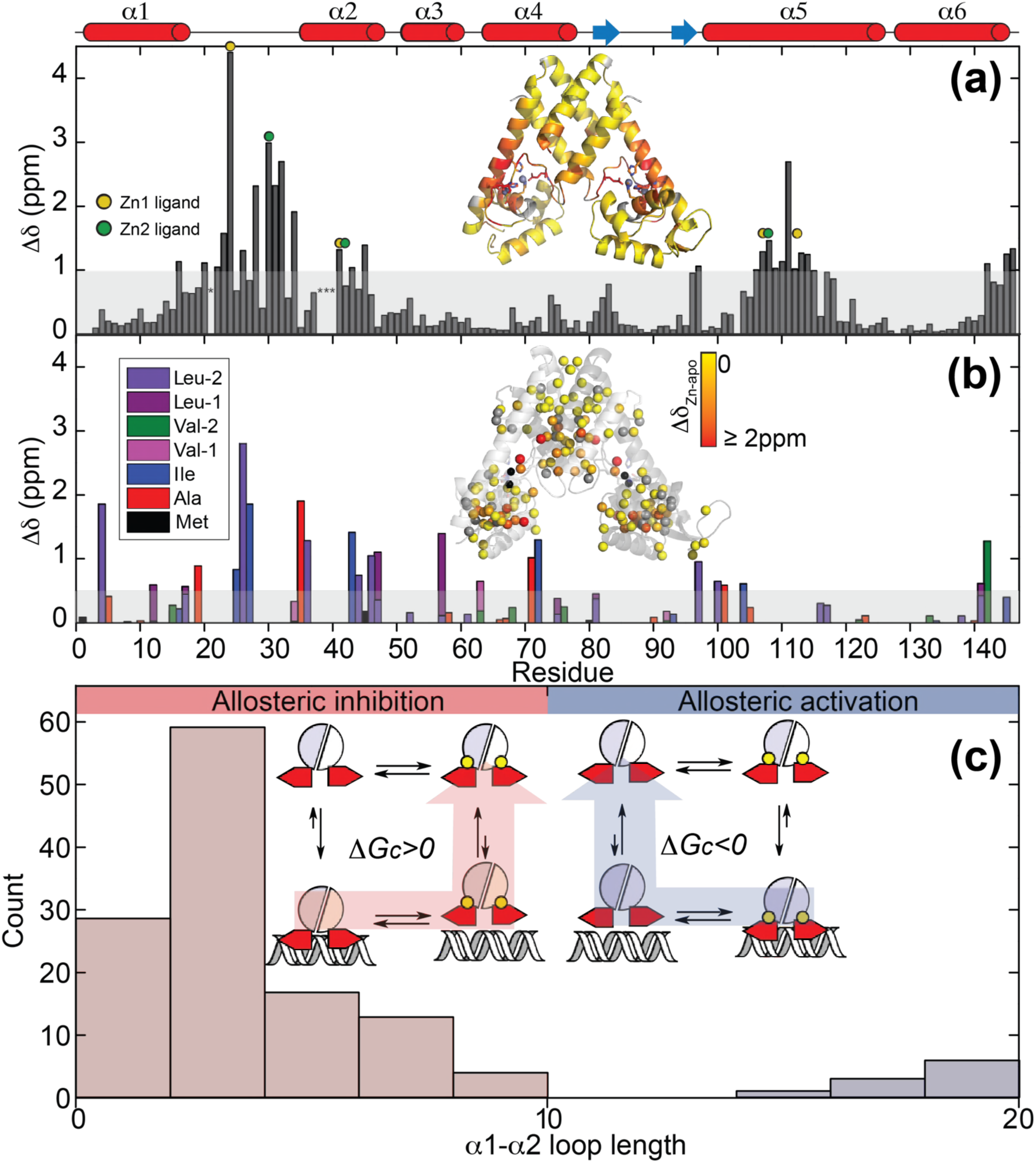
Chemical shift perturbation maps for Zn^II^ binding to AdcR for the backbone (a) and the sterospecifically assigned methyl groups (b) at pH 5.5, 50 mM NaCl, 35 °C. CSPs are painted on the ribbon representation of the structure of Zn^II^_2_ AdcR. The shaded bar in each case represents one standard deviation from the mean perturbation. Site 1 and site 2 ligands in the primary structure in panel (a) are denoted by the *yellow* and *green* circles, respectively; the asterisks at residue positions 21 and 38-40 indicate no assignment in the apo-state (see Materials and Methods). (c) Distribution of α1-α2 loops lengths in the reported structures in all MarR family of proteins (see Table S1 for an accounting of these structures). A postulated schematic representation of allosteric inhibition and activation are shown (*inset*), with shorter α1-α2 loops driving allosteric inhibition of the DNA binding upon ligand binding, and longer loops associated with allosteric activation (like that for AdcR/ZitR) upon ligand binding.

The changes in carbon chemical shifts in the central region of the α5 helix and the presence of strong NOEs to water for these residues are consistent with a kink in this helix in the apo-state (Fig. S3A-B), as is commonly found in other structurally characterized MarR repressors in DNA-binding inactive conformations (Zhu *et al.*, 2017b, Duval *et al.*, 2013). However, the kink is expected to be local and transient, since a TALOS+ analysis of chemical shifts predicts that the α5 helix remains the most probable secondary structure for all tripeptides containing these residues in the apo-state (Shen *et al.*, 2009) (Fig. S3C). The backbone changes in chemical shifts are accompanied by changes in the hydrophobic cores in the proximity of Zn^II^ binding as reported by the stereospecific sidechain methyl group chemical shift perturbation maps (Fig. 3B). Comparatively smaller perturbations extend to the α1 helix and the C-terminal region of the α6 helix, DNA-binding α4 helix (S74) and into the β-wing itself, consistent with a significant change in quaternary within the AdcR homodimer upon binding of both allosteric metal ions (Fig. 3A-B).

Overall, our NMR and SAXS data show that the main structural differences are localized in the region immediately surrounding the Zn^II^ coordination sites, giving rise to a change in the quaternary structure, while conserving the size and the overall secondary structure of the molecule. In particular, our data point to a kink in the α5 helix and a structural perturbation in the α1-α2 loop, which could be inducing a reorientation of the winged helix-turn-helix motifs relative to the dimerization domain.

In an effort to understand the functional consequences of the structural perturbations in the α1-α2 loop, we compared the length of the loop that connects the dimerization domain with the winged helical motif among AdcR and other members of the MarR family of known structure. This structural comparison and an extensive multiple sequence alignment reveals that only AdcR-like repressors harbor an α1-α2 loop larger than 10 residues (Fig. 3C). This loop extension does not seem to originate from an insertion, but from a change in secondary structure of the C-terminal region of the α1 helix (Fig. 3C). Moreover, in the Zn^II^ state that loop appears restricted by a hydrogen-bond network between the Zn^II^ binding site and the DNA binding domain (Chakravorty *et al.*, 2013). The Zn^II^-ZitR crystal structure similarly has an α1-α2 loop that is restricted by metal coordination chemistry and other intermolecular contacts with the dimerization and DNA binding domains, despite lacking an identifiable hydrogen-bond network (Zhu *et al.*, 2017c). Overall, our analysis suggests that the flexibility of this loop prevents DNA-binding, while the interactions formed in response to Zn^II^ coordination may be important in allosteric activation of DNA binding. Such a dynamical model contrasts sharply with a rigid body motion mechanism as previously suggested for other MarRs (Alekshun *et al.*, 2001, Chang *et al.*, 2010, Dolan *et al.*, 2011, Saridakis *et al.*, 2008, Birukou *et al.*, 2014, Radhakrishnan *et al.*, 2014), thus motivating efforts to understand how conformational dynamics impacts biological regulation by Zn^II^ in AdcR.

### Zn^II^-induced changes in AdcR conformational plasticity along the backbone

We therefore turned to an investigation of protein dynamics in AdcR. ^15^N*R*_1_, *R*_2_, and steady-state heteronuclear ^15^N{^1^H} NOEs provide information on internal mobility along the backbone, as well as on the overall protein tumbling rate (Fig. 4A-D; Fig. S4). The *R*_1_ and *R*_2_ data reveal that Zn^II^_2_ AdcR tumbles predominantly as a single globular unit in solution (Fig. 4B; Fig. S4) with a molecular correlation time (τ_c_) of 18.7 ± 0.1 ns, very similar to the τ_c_ value predicted for the dimer at 35 °C (18.9 ns in D_2_O). The β-wing region tumbles independently from the rest of the molecule (Fig. 4B). These data also reveal that the α1-α2 linker region that donates the E24 ligand to Zn^II^ binding site 1 is ordered to an extent similar to the rest of the molecule. In striking contrast, in apo-AdcR, the dimerization and DNA-binding domains have significantly smaller τ_c_ values (10.9 ± 0.5 ns, Fig. 4A), close to that expected if these domains tumble independently of one another in solution; in addition, the α1-α2 loop is highly dynamic in the apo-state (see also Fig. S4). These findings are consistent with the SAXS data, which show that apo-AdcR is less compact than the Zn^II^_2_ state. As in the Zn^II^_2_ state, the β-wing tumbles independently of the rest of the molecule, revealing that a change in the flexibility or orientation of the β-hairpin is likely not part of the allosteric mechanism, contrary to what has been proposed for other MarRs on the basis of crystal structures alone (Liu *et al.*, 2017, Deochand & Grove, 2017, Kim *et al.*, 2016). Overall, the ^15^N relaxation data for backbone amides show that Zn^II^ binding leads to a reduction of mobility of the α1-α2 loop, which in turn, decreases the dynamical independence the DNA-binding and dimerization domains, thereby stabilizing a conformation that tumbles in solution as a single globular unit.

**Figure 4.**
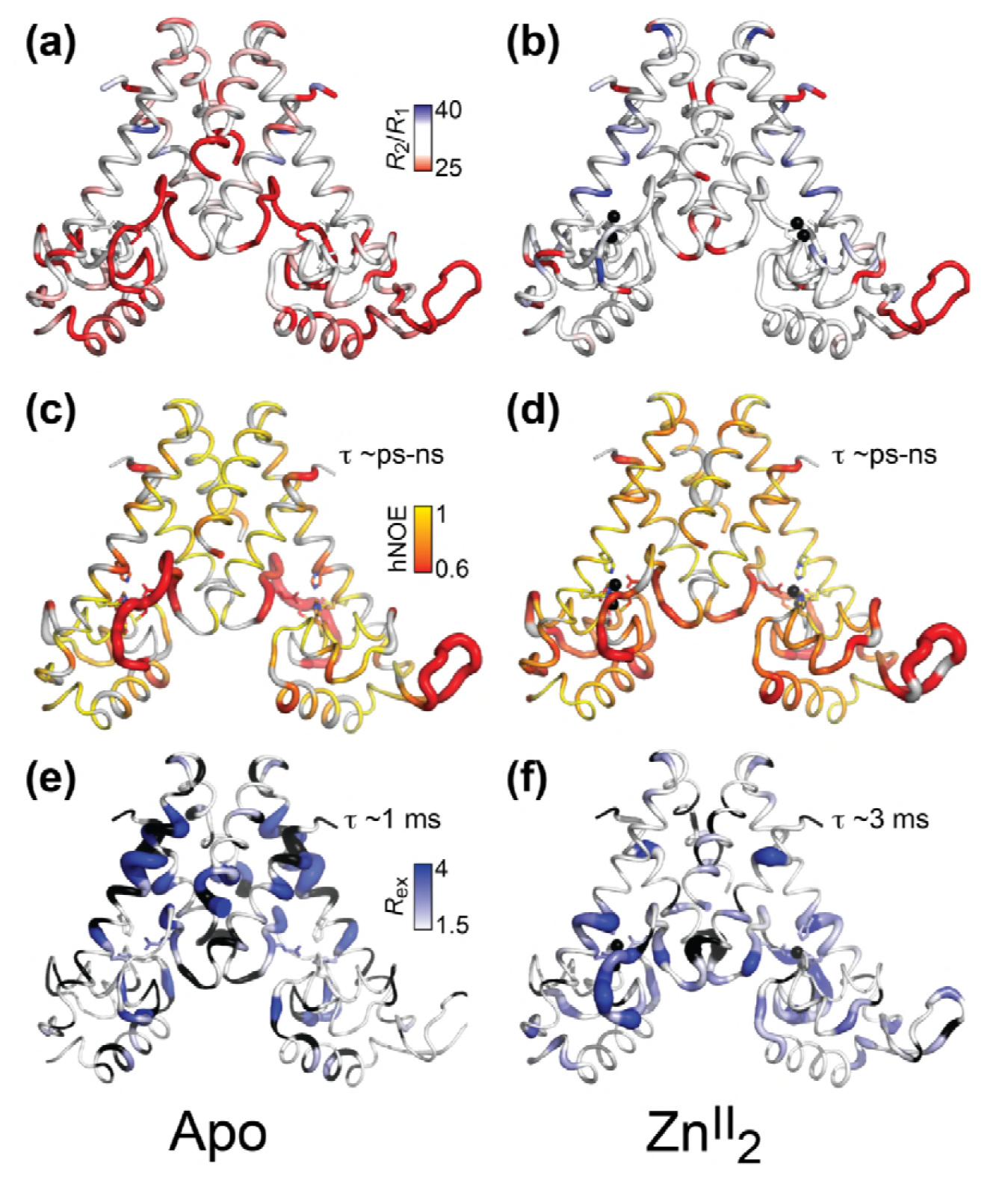
Dynamical characterization of the apo- (a) (c) (e) and Zn^II^_2_ (b) (d) (f) AdcR conformational states. Backbone ^1^H-^15^N amide *R*_2_/*R*_1_ for apo- (a) and Zn^II^_2_ AdcR (b) painted onto the 3tgn structure (Guerra *et al.*, 2011). Heteronuclear NOE analysis of apo- (c) and Zn^II^_2_ (d) AdcR with the values of the ^15^N-{^1^H}-NOE (hNOE) painted onto the 3tgn structure. Values of R_ex_ determined from HSQC ^15^N-^1^H CPMG relaxation dispersion experiments at a field of 600 MHz for the apo- (e) and Zn^II^_2_ (f) AdcRs (see Fig. S6 for complete data). Similar results were obtained at 800 MHz (Fig. S6). Zn^II^ ions are shown as black spheres and residues excluded due to overlapped are shown in gray and yellow. The width of the ribbon reflects the value represented in the color bar.

To further probe the reduction of flexibility upon Zn^II^ binding, we investigated sub-nanosecond backbone mobility as reported by the steady-state heteronuclear ^15^N{^1^H} NOEs (Fig. 4C-D). These hNOE data confirm that the internal mobility of the apo-state on this timescale mainly localizes to the α1-α2 loop and the central region of the α5 helix, around E107 (Zn^II^ site 2 ligand) and H108 and H112 (Zn^II^ site 1 ligands). The short-timescale flexibility in this region is significantly restricted upon Zn^II^ binding, but somewhat paradoxically leads to an *increase* in sub-nanosecond backbone motion in the DNA-binding domain, particularly in the α3 helix and the N-terminal region of the α4 helix, which harbors the key DNA-binding determinants (Fig. S1A) (Zhu *et al.*, 2017c). The quenching of sub-nanosecond mobility in the α1-α2 loop by Zn^II^ is accompanied by a corresponding increase in mobility on the μs-ms (slow) timescale in this region (Fig. 4F). In addition, the slow timescale backbone dynamics show a restriction of a conformational sampling in a band across the middle of the dimerization domain, including the upper region of the α5 helix, the N-terminus of α1, and the C-terminus of α6 (Fig. 4E-F). These slow motions in the apo-state likely report on a global breathing mode of the homodimer reflective of the conformational ensemble, which is substantially restricted upon Zn^II^ binding.

These large differences in structure and dynamics between the apo and Zn^II^_2_ AdcRs suggest an allosteric mechanism that relies on a redistribution of internal mobility in both fast- and slow timescale regimes, rather than one described by a rigid body motion. This mobility redistribution restricts the flexibility of the ligand binding site from the sub-nanosecond timescale in the apo-form to the millisecond timescale in the Zn^II^_2_ state (Fig. 4C,F). This restriction links the motion of the two functional domains (Fig. 4A-B) and locks AdcR in a triangular shape compatible with DNA binding. On the other hand, Zn^II^ enhances the internal flexibility in the DNA binding domain (Fig. 4C-D), which other studies show plays a role in sequence recognition and high affinity binding, particularly on the side chains (Capdevila *et al.*, 2017a, Kalodimos *et al.*, 2004, Anderson *et al.*, 2013).

### Zn^II^-induced perturbations of side chain conformational disorder in AdcR

Unlike the backbone, perturbations in side chain flexibility in the sub-nanosecond timescale are capable of reporting on the underlying thermodynamics of Zn^II^ binding and the role of conformational entropy (Δ*S*_conf_) in the allosteric mechanism. These perturbations potentially pinpoint residues with functional roles, *i.e.*, allosteric hotspots (Capdevila *et al.*, 2017a), with the change in the methyl group order parameter (Δ*S*^2^_axis_) upon ligand binding employed as a dynamical proxy (Capdevila *et al.*, 2017a, Caro *et al.*, 2017). Thus, if the motional redistribution observed in the backbone upon Zn^II^ binding is accompanied by changes in the dynamics on the side chains, particularly those in the DNA binding regions, these fast internal dynamics could affect the entropy of the metal binding and play a major role in the allosteric mechanism. To test these ideas, we first measured the axial order parameter, *S*^2^_axis_, for all 82 methyl groups, comparing the apo- and Zn-bound states of AdcR (Fig. 5A, Fig. S5). These dynamics changes are overall consistent with the stiffening observed along protein backbone, *e.g.*, in the α1-α2 loop; L26, in particular, is strongly impacted, changing motional regimes, |Δ*S*^2^_axis_|>0.2) (Frederick *et al.*, 2007). This stiffening prevails all over the molecule, leading to a small net decrease in conformational entropy upon Zn^II^ coordination (–*T*Δ*S*_conf_= 3.4 ± 0.4 kcal mol^−1^) (Fig. 5A). However, as has been previously shown for other transcriptional regulators (Capdevila *et al.*, 2017a, Tzeng & Kalodimos, 2012), the binding of the allosteric ligand Zn^II^ actually leads to a redistribution of sidechain mobility throughout the entire molecular scaffold. Interestingly, most of the methyl groups that change motional regimes are located in the DNA binding domain (Fig. S7). In particular, the side chain flexibility of many residues in the α3 helix *increases*, including L47, L57, L61, while a small hydrophobic core in the C-terminus of the α4 helix stiffens significantly, *e.g.*, L81, V34. These changes are accompanied by perturbations in the dynamics at the dimer interface, *i.e.*, L4, I16, V14, in both motional regimes as reported by Δ*S*^2^_axis_ and Δ*R*_ex_ (in the μs-ms timescale), the latter derived from relaxation dispersion experiments (Table S2; Fig. S5C).

**Figure 5.**
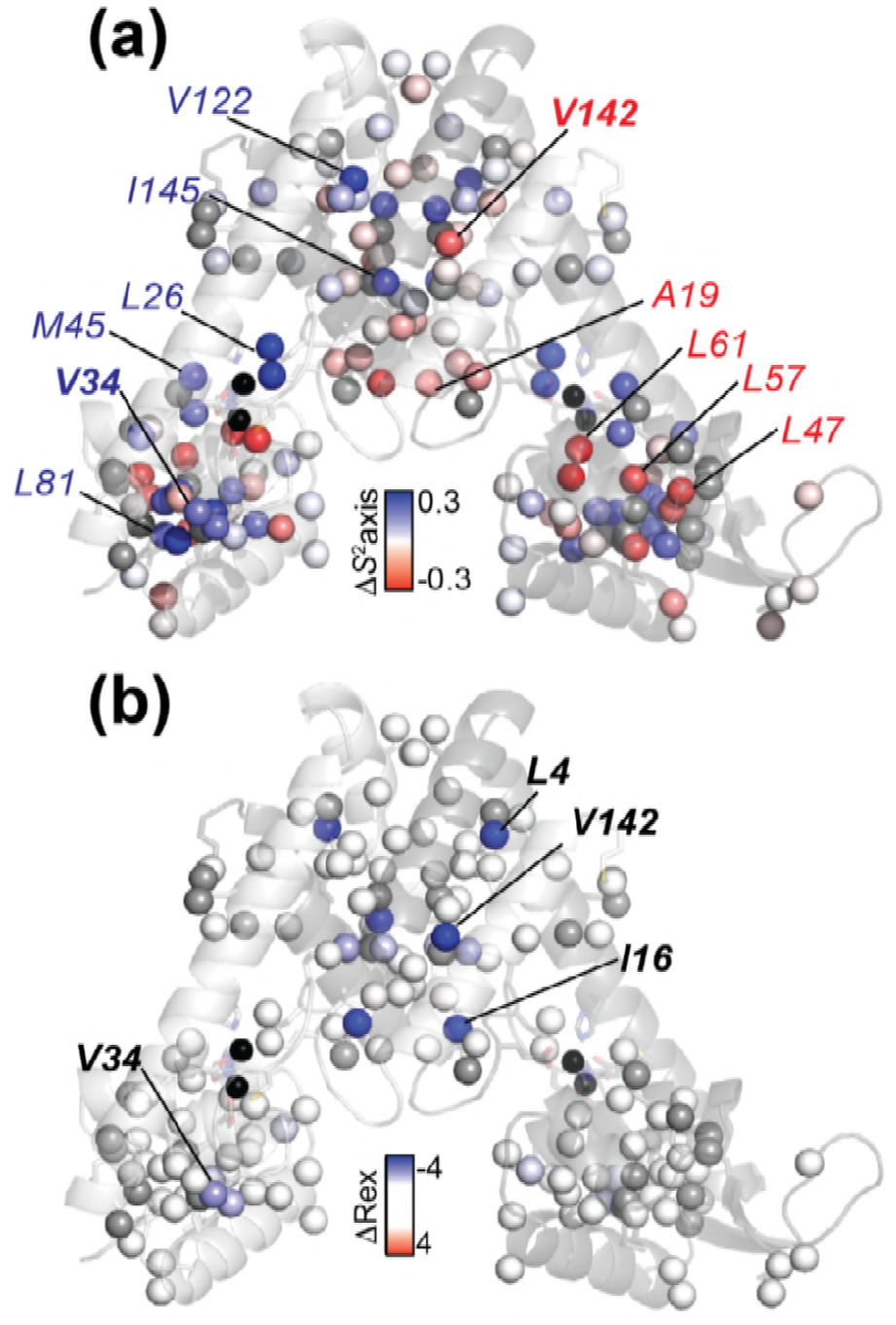
Effect of Zn^II^ binding to AdcR on the site-specific stereospecifically assigned methyl group axial order parameter, *S*^2^_axis_ (a) and *R*_ex_ (b) plotted as Δ*S*^2^_axis_ (*S*^2^_axis_^Zn^ – *S*^2^_axis_^apo^) and Δ*R*_ex_ (*R*_ex_^Zn^ – *R*_ex_^apo^) values, respectively, mapped onto the structure of Zn^II^_2_ AdcR (3tgn). A Δ*S*^2^_axis_ <0 indicates that the methyl group becomes *more* dynamic in the Zn^II^_2_-bound state, while Δ*R*_ex_<0 indicates quenching of motion on the μs-ms timescale in the in the Zn^II^_2_-bound state. See Fig. S5 for a graphical representation of all *S*^2^_axis_ and *R*_ex_ values in each conformation from which these differences were determined, and Fig. S6 for summary of all dynamical parameters measured here. Residues harboring methyl groups that show major dynamical perturbations on Zn^II^ binding are highlighted, with selected residues subjected to cavity mutagenesis (Fig. 6; Table S2).

**Figure 6.**
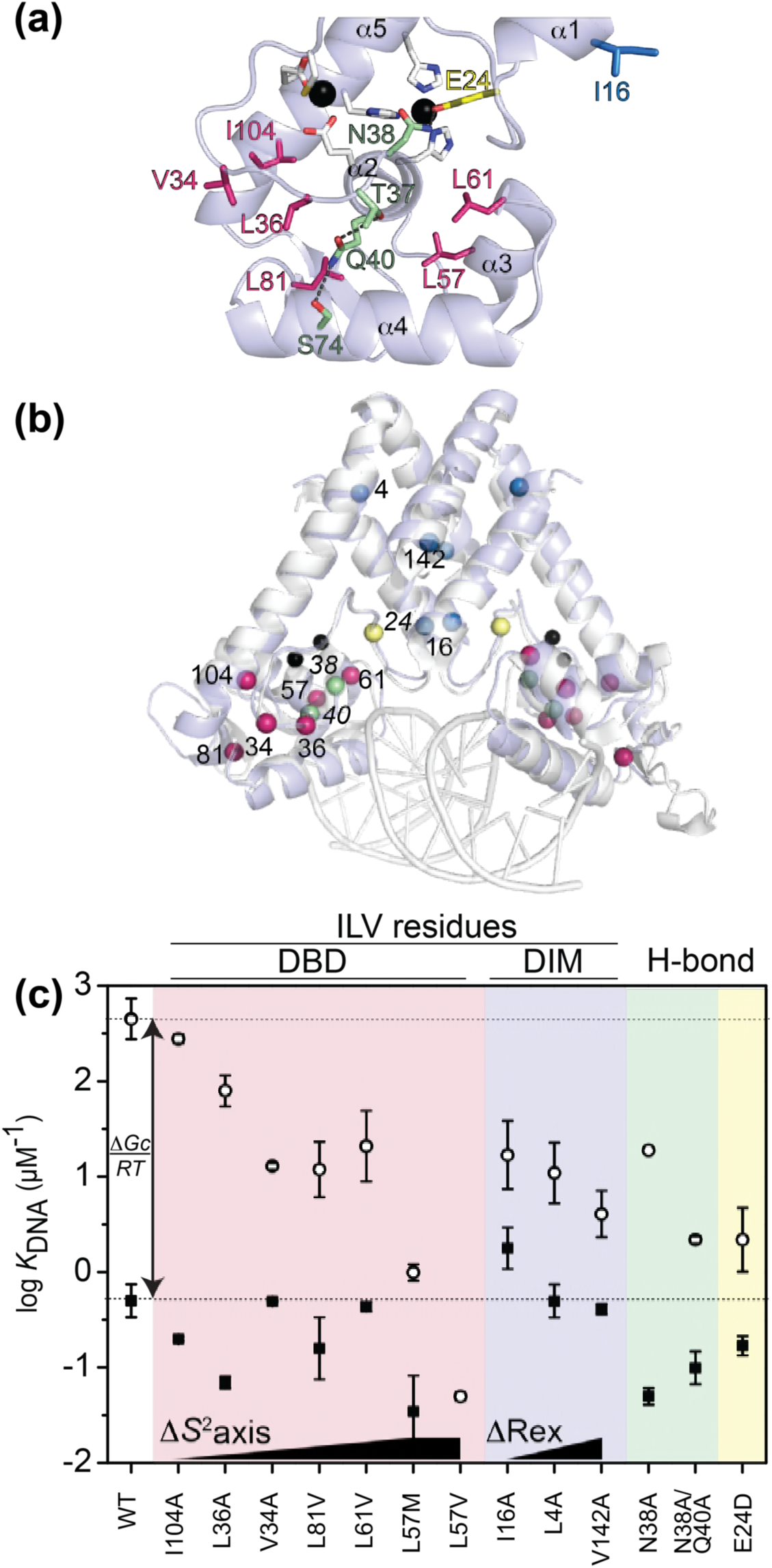
Graphical summary of the functional properties of AdcR cavity and hydrogen bonding mutants. (a) Zoom of the DNA binding domain (DBD) of one of the two Zn^II^_2_-bound AdcR protomers highlighting the residues targeted for mutagenesis (cavity mutants, *red stick*; hydrogen-bonding pathway mutants, *green stick*; zinc ligand E24, *yellow stick*), with the helical elements (α1-α5) indicated. (b) Cα positions of the residues targeted for cavity mutagenesis in the DNA binding domain (DBD) (*red spheres*) and in the dimerization domain (DIM) (*blue spheres*); other residues targeted for substitution in the hydrogen-bonding pathway (N38, Q40; *green spheres*) and zinc ligand E24 (*yellow spheres*) highlighted on the structure of the Zn^II^_2_ ZitR-DNA operator complex (Zhu *et al.*, 2017c); Zn^II^ ions (*black spheres*). (c) Coupling free energy analysis for all AdcR mutants highlighted using the same color scheme as in panels (a), (b). DBD, DNA-binding domain; DIM, dimerization domain; H-bond, hydrogen binding mutants. Lower horizontal line, *K*_DNA_ for wild-type apo-AdcR; upper horizontal line, *K*_DNA_ for wild-type Zn^II^_2_ AdcR, for reference. The trend in ΔS2axis and ΔRex is qualitatively indicated (see Table S2). These residues are conserved to various degrees in AdcR-like repressors (Fig. S12).

### On-pathway and off-pathway allosterically impaired mutants of AdcR

Our previous work (Capdevila *et al.*, 2017a) makes the prediction that “dynamically active” sidechains (methyl groups with |Δ*S*^2^_axis_|≥0.2 upon Zn^II^ binding) (see Fig. 5) are crucial for allosteric activation of DNA binding by Zn^II^. To test this prediction, we prepared and characterized several mutant AdcRs in an effort to disrupt allosteric activation of DNA binding, while maintaining the structure of the dimer and high affinity Zn^II^ binding. Since it was not clear *a priori* how mutations that perturb mobility distributions in one timescale or the other (sub-ns or μs-ms) would impact function, we focused on two kinds of substitution mutants: cavity mutants of dynamically “active” methyl-bearing side chains positioned in either the DNA binding or the dimerization subdomains (Fig. 6A, B) (Capdevila *et al.*, 2017a), and substitutions in the hydrogen-bonding pathway in the Zn-state that may contribute to the rigidity of the α1-α2 loop in Zn^II^_2_-AdcR (Fig. 6A) (Chakravorty *et al.*, 2013). We measured DNA binding affinities of the apo and zinc-saturated Zn^II^_2_-states, and calculated the allosteric coupling free energy, Δ*G*_c_, from Δ*G*_c_=–*RT*ln(*K*_Zn,DNA_/*K*_apo, DNA_) (Giedroc & Arunkumar, 2007) (Fig. 6C and Table S2). All mutants are homodimers by size-exclusion chromatography (Fig. S9) and all bind the first equivalent of Zn^II^ tightly as wild-type AdcR (Fig. S10, Table S3).

*DNA-binding domain mutants.* The redistribution fast time scale side-chain dynamics in the DNA binding domain is delocalized throughout the different secondary structure motifs. Thus, we prepared several cavity mutants of methyl-bearing residues in the α3 (L57, L61), α4 (L81) and α5 (I104) helices, as well as two residues in the α1-α2 loop in close proximity to the N-terminus of α2, V34 and L36. I104 is the most distal from the bound DNA in the Zn^II^_2_ ZitR-DNA complex (Zhu *et al.*, 2017c), and is not dynamically active in AdcR (|Δ*S*^2^_axis_|<0.1; Δ*R*_ex_<1.0); thus, the I104A mutant is predicted to function as a control substitution. V34 and L36 are dynamically active on both timescales, which is not surprising since the α1-α2 loop folds upon Zn^II^ binding to AdcR (*vide supra*) (Zhu *et al.*, 2017c). In contrast, L57, L61 and L81 are characterized by significant perturbations in Δ*S*^2^_axis_ only (|Δ*S*^2^_axis_|≥0.2), with L81 stiffening and L57 and L61 methyls in the α3 helix becoming significantly more dynamic upon Zn^II^ binding (Fig. 5A, Table S2).

As expected, I104A AdcR is characterized by DNA binding affinities in the apo- and Zn-states just ≈2-fold lower than wild-type AdcR, returning a Δ*G*_c_ that is not statistically different from wild-type AdcR (Fig. 6C). Functional characterization of all other cavity mutants in the DNA binding domain results in a ≈5-10-fold decrease or greater (L57V AdcR; Table S2) in the DNA binding affinity of the apo-state (Fig. 6C), with Zn^II^ binding inducing markedly variable degrees of allosteric activation (Fig. 6C). L36A, closest to the α2 N-terminus, is most like wild-type AdcR, while V34A AdcR is severely crippled in allostery, with *K*_Zn,DNA_ some 200-fold lower than wild-type AdcR, and Δ*G*_c_ ≈2-fold lower, from –4.0 to –2.2 kcal mol^−1^. L81V and L61A AdcRs are comparably perturbed, and L57M AdcR even more so (Δ*G*_c_≈–2.0 kcal mol^−1^).

We emphasize that these methyl-bearing side chains targeted for substitution are ≥95% buried and none are expected to be in direct contact with the DNA (Fig. 6B, Table SI). These data provide strong support for the idea that those methyl-bearing side chains in the DNA-binding domain that exhibit large changes in conformational entropy (as measured by Δ*S*^2^_axis_) make significant contributions to both DNA binding and allosteric activation by Zn^II^. This result highlights the contribution that dynamical redistribution within the DNA-binding domain makes for AdcR function, as has been observed in other transcriptional regulators (Tzeng & Kalodimos, 2012; Capdevila *et al*., 2017a).

*Hydrogen*-*bonding mutants.* A hydrogen-bonding pathway in AdcR (Chakravorty *et al.*, 2013) has previously been proposed to transmit the Zn^II^_2_ binding signal to the DNA binding domain. In this pathway, the Oε1 atom from the Zn^II^ ligand E24 accepts a hydrogen bond from the carboxamide side chain of N38. N38 is the +1 residue of the α2 helix, which is then connected to the α4 helix via a hydrogen bond between the Q40 and S74 side chains; further, Q40 accepts a hydrogen bond from the γ-OH of T37 as part of a non-canonical helix N-capping interaction (Guerra *et al.*, 2011) (Fig. 6A). We expect that regardless of the impact that these interactions have on the overall energetics of Zn^II^ binding, they are important in the restriction of fast-time scale dynamics in the α1-α2 loop. We therefore targeted residues E24 (Zn-ligand and H-bound acceptor), N38 and Q40, by characterizing two single mutants, E24D and N38A, and the double mutant, N38A/Q40A AdcR. Although all three mutants undergo allosteric switching as revealed by ^1^H—^15^N TROSY spectra (Fig. S11), as with all other DNA-binding domain mutants, all three exhibit ≈5-10-fold decreases in apo-state DNA-binding affinity (Fig. 6C; Table S2). While the single mutant N38A binds Zn^II^ to give Δ*G*_c_ of ≈–3.5 kcal mol^−1^, quite similar to that of wild-type AdcR, in marked contrast, N38A/Q40A AdcR is functionally perturbed, characterized by a Δ*G*_c_ of ≈–1.9 kcal mol^−1^ and is E24D AdcR that target a Zn^II^ binding residue (Fig. 6C). These perturbations provide additional evidence that this hydrogen-bonding pathway may contribute to the motional restriction of the α1-α2 loop, jointly with a redistribution of internal dynamics in the DNA binding domain. This effect can be perturbed directly by mutation of “dynamically active” sidechains (L81V, L61V, L57M) or by significantly impacting the interactions that restrict the loop (N38A/Q40A).

*Dimerization domain mutants.* To test the functional role of the dimerization domain in dynamical changes, we targeted three methyl-bearing residues in this domain, including L4 and I16 on opposite ends of the α1 helix and V142, near the C-terminus of the α6 helix (Fig. 6B). L16 is closest to the intervening minor groove of the DNA operator, while V142 and L4 are increasingly distant from the DNA. These residues are primarily active in slow timescale dynamics, with Zn^II^-binding quenching side chain mobility on the μs-ms timescale, *i.e.*, global motions, but relatively smaller changes in Δ*S*^2^_axis_ (Fig. 5B; Table S2). Cavity mutants of these residues (I16A, L4A and V142A) bind DNA in the apo-state with wild-type like affinities, but each is allosterically strongly perturbed, with only ≈10-20-fold allosteric activation by Zn^II^, giving Δ*G*_c_ values of –1.4 to –1.8 kcal mol^−1^.

These findings suggest that Zn^II^-dependent quenching of global motions far from the DNA binding domain play a significant role in allostery in this system. Our characterization of allosterically compromised mutants that affect site-specific conformational entropy (L81V, L61V, L57M) and conformational exchange (V34A, L4A, I16A) provides evidence for two classes of functional dynamics in AdcR that comprise different regions of the molecule, operating on different timescales (from sub-nanoseconds to milliseconds). Thus, we propose that a Zn^II^-dependent redistribution of internal dynamics quenches global, slow motions in the dimer, yet enhances local dynamical disorder in the DNA binding domain, which can ultimately be harnessed to maximize contacts at the protein-DNA interface.

## Conclusions

Members of the multiple antibiotic resistance repressor (MarR) family of proteins comprise at least 12,000 members (Capdevila *et al.*, 2017b), and many have been subjected to significant structural inquiry since the original discovery of the *E. coli mar* operon and characterization of *E. coli* MarR some 25 years ago (Cohen *et al.*, 1993, Seoane & Levy, 1995). The crystallographic structure of this prototypical *E. coli* MarR appeared a number of years later (Alekshun *et al.*, 2001) and has inspired considerable efforts to understand the inducer specificity and mechanisms of transcriptional regulation in *E. coli* MarR (Hao *et al.*, 2014) and other MarR family repressors (Grove, 2013), which collectively respond to an wide range of stimuli, including small molecules, metal ions, antibiotic and oxidative stress (Deochand & Grove, 2017). We have examined the wealth of the crystallographic data available from 137 MarR family repressor structures solved in a variety of functional states, including DNA-binding competent, DNA-binding incompetent and DNA-bound states (Fig. 1). This analysis of the crystal structures suggests that conformational selection induced by ligand binding or thiol oxidation must be operative in a significant number of these repressor systems. Here, we present the first site-specific dynamics analysis of any MarR family repressor in solution, and establish that conformational dynamics on a range of timescales is a central feature of Zn^II^-dependent allosteric activation of DNA operator binding by the zinc uptake regulator, *S. pneumoniae* AdcR (Reyes-Caballero *et al.*, 2010) and closely related repressors (Zhu *et al.*, 2017c).

We explored dynamics in the sub-nanosecond and μs-ms timescales with residue-specific resolution, both along the backbone, as measured by N-H bond vectors, and in the methyl groups of the methyl-bearing side chains of Ala, Met, Val, Leu and Ile. These measurements, coupled with small angle x-ray scattering measurements of both conformational states, lead to a consistent picture of allosteric activation by Zn^II^ in AdcR. The apo-state conformational ensemble is far broader than the Zn^II^_2_ state, and features dynamical uncoupling of the core DNA-binding and dimerization domains, facilitated by rapid, low amplitude motions in the α1-α2 loop and the α5 helix in the immediate vicinity of the Zn^II^ coordinating residues. This motion is superimposed on a much slower, larger amplitude mobility across the dimerization domain, far from the DNA interface, affecting both backbone amide and side chain methyl groups (Fig. 4-5). Zn^II^ binding substantially quenches both the low amplitude internal motions and global, larger amplitude movements, while driving a striking redistribution of these dynamics into the DNA-binding domain.

As we observed previously for another Zn^II^ metalloregulatory protein, Zn^II^ binding induces a net global conformational stiffening, superimposed on pockets of increased dynamical disorder, particularly in the α3-α4 region of the DNA binding domain (Fig. 5A). It is interesting to note that the structures of Zn^II^_2_-bound AdcR and DNA-bound Zn^II^_2_ ZitR differ most strongly in the α3-α4 region (Fig. 6B), suggesting that internal dynamics in this region may be functionally important in enhancing DNA binding affinity. To test the functional importance of fast-time scale motions in the DNA binding domain, we exploited the side chain dynamics results (Fig. 5) (Capdevila *et al.*, 2017a) to design cavity substitutions of dynamically active residues (Fig. 6). We generally find that cavity substitutions in the DNA binding domain are strongly deleterious for residues that are dynamically active in the fast timescale (|Δ*S*^2^_axis_|>0.2), *i.e.* L81, L61, L57. These findings confirm a functional role of these changes in dynamics (Capdevila *et al.*, 2017a) and suggest that Zn^II^_2_-bound AdcR has an optimal distribution of internal dynamics that if perturbed, leads to weakened DNA binding.

Crystallographic studies suggest that DNA binding in MarR repressors is optimized by precisely tuning interactions with the DNA operator sequence, resulting in a favorable Δ*H* of binding (Hong *et al.*, 2005, Dolan *et al.*, 2011, Quade *et al.*, 2012, Birukou *et al.*, 2014, Zhu *et al.*, 2017a, Gao *et al.*, 2017, Otani *et al.*, 2016) with the functional consequences of ligand binding known to vary widely among individual MarRs (Deochand & Grove, 2017). We propose here, based on this and previous work, that tighter DNA binding can be achieved by optimization of side-chain dynamics that give rise to a more favorable conformational entropy term (Δ*S*_conf_) and the functional consequences of ligand binding can be predicted based on the protein internal motions. We show here that single point mutations in AdcR, sufficient to impact internal motions, result in destabilization of the ternary complex. We showed previously that ligand binding can inhibit formation of the DNA complex by restricting the coupled fast motions and concerted slower motions that contribute to a favorable conformational entropy of DNA binding (Capdevila *et al.*, 2017a). This can potentially be the case also for introduction of dynamic elements, *i.e.*, loops or disordered regions (Pabis *et al.*, 2018, Campbell *et al.*, 2016). Thus, in the context of evolution of the MarR repressors, we propose that two allosteric modes, activation and inhibition, may have evolved by tuning the conformational entropy contribution to DNA binding (Fig. 7).

**Figure 7.**
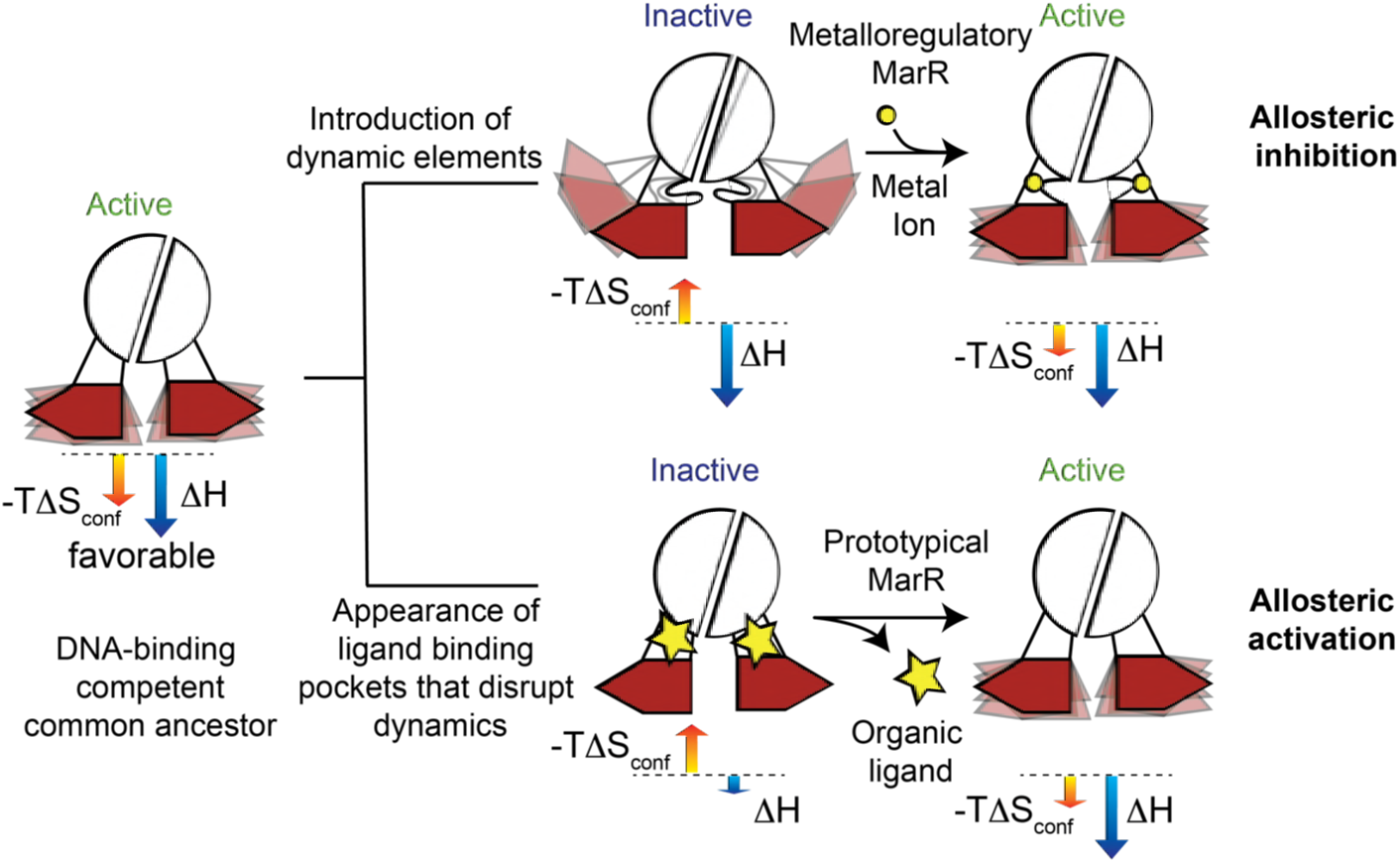
Entropically driven model for how conformational dynamics can be harnessed to evolve allosteric activation (*upper right*) vs. allosteric inhibition (*lower right*) in the same molecular scaffold. This model suggests that dynamic properties of the active states have been conserved to give rise to a more favorable conformational entropy. In the metalloregulatory MarRs (AdcR, ZitR), the inactive state shows perturbed dynamics over a range of timescales; apo-AdcR therefore exhibits low affinity for DNA. Metal ion (*yellow circle*) coordination quenches both local and global modes in the dimerization domain and linkers, while inducing conformational disorder in the DNA-binding domain that enhances DNA binding affinity, thus stabilizing a conformation that has high affinity for DNA and giving rise to a favorable conformational entropy. For prototypical MarRs, where the ligand (*yellow star*) is an allosteric inhibitor, ligand binding narrows the conformational ensemble to a DNA-binding “inactive” conformation decreasing the enthalpic contribution to DNA binding, while perturbing fast time scale dynamics that give rise to an unfavorable conformational entropy for DNA-binding.

Allosteric inhibition could have arisen by evolution of a ligand binding pocket where inducer recognition disrupts internal dynamics and increases the conformational entropic cost of binding to DNA, as we have previously shown for an ArsR family protein (Capdevila *et al.*, 2017a). Although this hypothesis has not been tested experimentally on any MarR, molecular dynamics simulations show that DNA binding-impaired mutants of MexR differ from the wild-type repressor in the nature of the dynamical connection between the dimerization and DNA binding domains (Anandapadamanaban *et al.*, 2016). This dynamical connectivity is in fact, exploited by the binding the ArmR peptide, leading to DNA dissociation (Anandapadamanaban *et al.*, 2016, Wilke *et al.*, 2008). We propose that conformational entropy can contribute to other mechanisms of allosteric inhibition to yield a repressor that binds tightly to the operator sequence and yet has the ability to readily evolve new inducer specificities.

On the other hand, allosteric activation could have evolved by perturbing internal dynamics on the apo-protein and increasing the conformational entropic cost of DNA binding. This perturbation could arise by the introduction of loops or disordered regions in the apo-protein that could be compensated by a ligand binding event that restores the internal dynamics to yield a more favorable entropy contribution to DNA binding (Fig. 7). Similarly, it has been shown that another well-studied, allosterically activated, bacterial regulator, catabolite activator protein (CAP) (Tzeng & Kalodimos, 2013, Tzeng & Kalodimos, 2012) is able to harness conformational entropy to increase DNA binding affinity upon ligand binding. In the case of MarRs, the far longer α1-α2 linker in AdcRs (Fig. 3) may have been an important intermediate determinant in the evolution of allostery in AdcR, given the key role this loop plays in dynamical uncoupling of the dimerization and DNA-binding domains in the ligand-free state (Fig. 7).

This model makes the prediction that if conformational entropy can be harnessed to bind DNA with high affinity, perturbations introduced by ligand binding or subtle change in protein sequence that conserve the molecular scaffold can easily lead to inactivation of DNA binding (Fig. 7). It is interesting to note that mutations that lead to inactivation are not necessarily part of a physical pathway with the DNA binding site (Clarke *et al.*, 2016), since they only need to affect dynamical properties that are likely delocalized in an extended network. Notably, single point mutants in the dimerization domain of various MarR family repressors have been shown to modulate allostery and DNA binding (Anandapadamanaban *et al.*, 2016, Deochand *et al.*, 2016, Liguori *et al.*, 2016, Duval *et al.*, 2013, Alekshun & Levy, 1999, Andresen *et al.*, 2010). In the case of AdcR, structural perturbations induced by Zn^II^ binding are essentially confined to the Zn^II^ binding pocket, *i.e.*, the α1-α2 loop and the α5 helix proximal to the Zn^II^ donor ligands (Fig. 3).

In striking contrast, dynamical perturbations extend all over the molecule, and feature many residues that are far from either ligand binding site, and are dynamically active on the sub-nanosecond and/or μs-ms timescales (Figs. 4-5). Thus, the conformational entropy contribution being inherently delocalized and easily perturbed can enable rapid optimization of new inactivation mechanisms that would allow new biological functionalities to arise (Fig 7). Moreover, we suggest that changes in the site-specific dynamics, derived from differences in the amino acid sequence, could evolve allosteric activation from allosteric inhibition in the context of the same overall molecular scaffold. These findings inspire efforts to explore the evolution of allostery in this remarkable family of transcriptional repressors, by exploiting an allosterically crippled AdcR (see Fig. 6) to re-evolve functional allostery on this system.

## Materials and Methods

### AdcR mutant plasmid production

An overexpression plasmid for *S. pneumoniae* AdcR in a pET3a vector was obtained as previously described and was used as a template for the production of all mutant plasmids (Reyes-Caballero *et al.*, 2010). Mutant AdcR plasmids were constructed by PCR-based site-directed mutagenesis, and verified using DNA sequencing.

### Protein production and purification

AdcR plasmids were transformed into either *E. coli* BL21(DE3) pLysS or Rosetta cells. *E. coli* cultures were either grown in LB media or M9 minimal media supplemented with ^15^NH_4_Cl as the sole nitrogen source with simple ^1^H,^15^N HSQC spectroscopy to assess the structural integrity of selected mutant proteins. Protein samples for backbone and methyl group assignments of AdcR were isotopically labeled using published procedures as described in our previous work (Capdevila *et al.*, 2017a, Arunkumar *et al.*, 2007), with all isotopes for NMR experiments purchased from Cambridge Isotope Laboratories. Protein expression and purification were carried out essentially as previously described (Reyes-Caballero *et al.*, 2010). All proteins were confirmed to have <0.05 molar equivalents of Zn(II) as measured by atomic absorption spectroscopy and were dimeric by gel filtration chromatography. The AdcR protein concentration was measured using the estimated molar extinction coefficient at 280 nm of 2980 M^−1^ cm^−1^.

### Small angle x-ray scattering experiments

Small angle and wide angle x-ray scattering data of the apo and Zn^II^_2_ states of AdcR was collected at three different protein concentrations (5 mg/mL, 2.5 mg/mL and 1.25 mg/mL) in buffer 25 mM MES pH 5.5, 400 mM NaCl, 2 mM EDTA/10 μM ZnCl_2_, 2 mM TCEP at sector 12ID-B at the Advanced Photo Source (APS) at Argonne National Laboratory. For each protein concentration and matching background buffer, 30 images were collected and averaged using NCI-SAXS program package. The scattering profile at each concentration was manually adjusted with the scale factor to remove the effect of concentration prior to subtraction of the scattering profile of the buffer. Scattering profiles of each protein concentration were then merged for further analysis. The GUINIER region was plotted with ln (I(*q*)) vs *q^2^* to check for monodispersity of the sample and to obtain *I_0_* and the radius of gyration (*R_g_*) within the range of *q_max_*^∗^*R_g_* <1.3. The *R_g_* values obtained for apo-AdcR and Zn(II)-bound-AdcR are 25.5 ± 0.9Å and 23.7 ± 1.1 Å, respectively. The scattering profiles of each AdcR conformational state was then normalized with *I*_0_. The compaction of each states of AdcR was examined using the Kratky plot for *q*<0.3 Å^−1^. Scattering profiles for apo and Zn^II^_2_ states of AdcR were then Fourier-transformed using GNOM of the ATSAS package to obtain the normalized pair-wise distance distribution graph (PDDF).

*Ab initio* modeling was performed using the program DAMMIF in a slow mode (Franke & Svergun, 2009). For each conformational state of AdcR, 10 models were obtained. These models were compared, aligned and averaged using the DAMSEL, DAMSUP, DAMAVER, DAMFILT, respectively, as described in the ATSAS package (http://www.embl-hamburg.de/bioSAXS). Normalized spatial discrepancy (NSD) between each pair of the models was computed. The model with the lowest NSD value was selected as the reference against which the other models were superimposed. Outliner models (2 models) with an NSD above mean + 2^∗^standard deviation of NSD were removed before averaging. For refinement, the averaged envelope of the first run was used as search volume for the second round of modeling. Modeling of the envelope of apo-AdcR was restrained by enforcing *P*_2_ rotational symmetry while that Zn^II^_2_ AdcR was restrained using compact, hallow and no-penalty constraints. Scattering profiles of crystal structures were calculated using the fast x-ray scattering (FOXS) webserver (https://modbase.compbio.ucsf.edu/foxs/) (Schneidman-Duhovny *et al.*, 2010).

### Mag-fura-2 competition assays

All mag-fura-2 competition experiments were performed on an ISS PC1 spectrofluorometer in operating steady-state mode or a HP8453 UV-Vis spectrophotometer as described in our previous work (Capdevila *et al.*, 2017a, Campanello *et al.*, 2013) using the following solution conditions: 10 mM Hepes, pH 7.2, 400 mM NaCl that was chelex treated to remove contaminating metals. 10 μM protein concentration was used for all and MF2 concentration ranged from 13-16 μM. These data were fit using a competitive binding model with DynaFit (Kuzmic, 1996) to determine zinc binding affinities for wild-type and each mutant AdcR using a four-site-nondissociable homodimer binding model, as previously described (Reyes-Caballero *et al.*, 2010) with *K_Zn_*= 4.9 × 10^6^ M^−1^ for mag-fura-2 fixed in these fits. *K*_1_ and *K*_2_ correspond to filling the two high affinity sites (site 1), and only a lower limits (≥10^9^ M^−1^) could be obtained for these sites; *K*_3_ and *K*_4_ were allowed to vary in the fit, and are reported in Table S3. Experiments were conducted 3 times for each AdcR variant. Errors of the binding constant parameters were estimated from global fits.

### NMR spectroscopy

NMR spectra were acquired on a Varian VNMRS 600 or 800 MHz spectrometer, each equipped with a cryogenic probe, at the Indiana University METACyt Biomolecular NMR laboratory. The two-dimensional spectra were processed using NMRPipe (Delaglio *et al.*, 1995). The three-dimensional spectra were acquired using Poisson-gap non-uniform sampling and reconstructed using hmsIST (Hyberts *et al.*, 2012) and analyzed using SPARKY (Goddard & Kneller) or CARA (http://cara.nmr.ch). Typical solution conditions were ∼500 μM protein (protomer), 25 mM MES pH 5.5, 50 mM NaCl, 1 mM TCEP, 0.02% (w/v) NaN_3_, and 10 % D_2_O. Some spectra were recorded at pH 6.0 as indicated. Our previous NMR studies of AdcR (Guerra *et al.*, 2011, Guerra & Giedroc, 2014) were carried out with samples containing ≈70% random fractional deuteration, pH 6.0, 50 mM NaCl, 35 °C; under those conditions, the backbone amides of residues 21-26 in the α1-α2 loop and harboring zinc ligand E24 as well as the N-terminal region of the α2 helix (residues 37-40) exhibited significant conformational exchange broadening in the apo-state and could not be assigned (Guerra *et al.*, 2011). In this work, we acquired comprehensive ^1^H-^15^N TROSY-edited NMR data sets at 600 and 800 MHz for a 100% deuterated AdcR sample in both apo- and Zn_2_-bound states at pH 5.5, 50 mM NaCl, 35° C. Under these conditions, only four backbone amides residues in the apo-state were broadened beyond detection (residues 21, 38-40); all were visible and therefore assignable in the Zn^II^_2_ state. Thus, the N-terminus of the α2 helix, including N38 and Q40 are clearly exchange broadened in the apo-state. Sidechains were assigned following published procedures as described in our previous work (Capdevila *et al.*, 2017a, Arunkumar *et al.*, 2007). The Leu and Val methyl resonances were distinguished using through-bond information such as HMCMCBCA or HMCM[CG]CBCA experiments (Tugarinov & Kay, 2003) which correlate the Leu or Val methyl resonances with other side chain carbon resonances. All apo-protein samples contained 1 mM EDTA. All Zn^II^_2_ samples contained 2 monomer mol equiv of Zn(II). Chemical shifts were referenced to 2,2-dimethyl-2-silapentane-5-sulfonic acid (DSS; Sigma) (Wishart & Sykes, 1994).

*S*^2^_axis_ of the Ile δ1, Leu δ1/δ2, Val γ1/γ2, Ala β, and Met ε methyl groups in apo and Zn(II)_2_ states were determined using ^1^H spin-based relaxation experiments at 600 MHz at 35.0 °C (Tugarinov *et al.*, 2007). *S*^2^_axis_ values, cross-correlated relaxation rates, η, between pairs of ^1^H-^1^H vectors in ^13^CH_3_ methyl groups were measured using Eq. 2

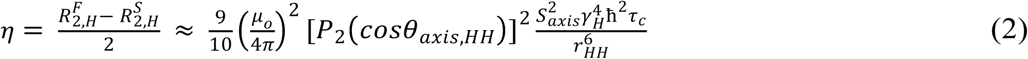

where τ_c_ is the tumbling time of the protein; *R*^F^_2,H_ and *R*^S^_2,H_ are the fast and slow relaxing magnetization, respectively; γ_H_ is the gyromagnetic ratio of the proton; and r_HH_ is the distance between pairs of methyl protons.

In order to obtain an approximation of the differences in fast and slow relaxation rates (2η, we measured the time-dependence of the cross peak intensities in a correlated pair of single and double quantum (2Q) experiments (Tugarinov *et al.*, 2007). Using various delay time, *T*, values (3, 5, 8, 12, 17, 22, and 27 ms, recorded in an interleaved manner), the rates of η were obtained by fitting ratios of peak intensities measured in pairs of experiments (*I*_a_ and *I*_b_, spin-forbidden and spin-allowed, respectively) with Eq. 3:

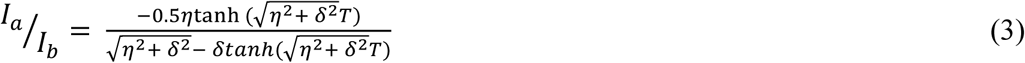

where *T* is the variable delay time, δ is a parameter that is related to the ^1^H spin density around the methyl group, and I_a_ and *I_b_* are the time dependencies of differences and sums, respectively, of magnetization derived from methyl ^1^H single-quantum transitions, as described (Tugarinov *et al.*, 2007). Peak heights and spectral noise were measured in Sparky (Lee *et al.*, 2015). A python script was used to fit the peak height ratios to η values and to determine *S*^2^_axis_ values in the apo- or Zn-bound states, as described previously (Tugarinov & Kay, 2004, Tugarinov *et al.*, 2007, Capdevila *et al*., 2017a). τ_c_ was obtained from Monte Carlo simulations with tensor2 software (Dosset *et al*., 2000), using *T*_1_, *T*_2_, and heteronuclear NOE (hNOE) recorded at 35 °C at 800 MHz, in 10% D_2_O. To enable a direct comparison of each AdcR allosteric state while overcoming the difficulty of determining an isotropic τ_c_ from tensor2 for apo-AdcR (which harbors dynamically independent domains), the measured τ_c_ for each state was obtained by adjusting *S*^2^_axis_ of alanine methyls to 0.85 since these methyl groups are generally motionally restricted in proteins (Igumenova *et al.*, 2006). For apo- and Zn^II^_2_ AdcRs, the τ_c_ obtained in this way is 18.9 ± 0.1 ns and 20.7 ± 0.1 ns respectively.

Relaxation dispersion measurements were acquired using a TROSY adaptation of ^15^N and a ^1^H-^13^C HMQC-based Carr–Purcell–Meiboom–Gill (CPMG) pulse sequence for amides from the backbone (Tollinger *et al.*, 2001) and methyl groups from the sidechains (Korzhnev *et al.*, 2004), respectively. Experiments were performed at 35 °C at 600 and 800 MHz ^1^H frequencies using constant time interval *T*=40 ms with CPMG field strengths (ν_CPMG_) of 50, 100, 150, 200, 250, 300, 350, 400, 450, 500, 600, 700, 850, and 1,000 Hz. Data were fitted to the two-site fast exchange limit equation, as discussed previously (Capdevila *et al.*, 2017a). These experiments were performed on duplicate at the two field strengths.

### DNA binding experiments and determination of allosteric coupling free energies (Δ*G_c_*)

For all DNA binding experiments a 28 bp double stranded DNA was obtained as previously described (Reyes-Caballero *et al.*, 2010) with the following sequence of the AdcO: 5’-TGATATAA<u>TTAAC</u>TG<u>GTAAA</u> CAAA ATGT[F]-3’. Apo AdcR binding experiments were conducted in solution conditions of 10 mM HEPES, pH 7.0, 0.23 M NaCl, 1 mM TCEP (chelexed), 10 nM DNA, 25.0 °C with 2.0 mM EDTA (for apo-AdcR) or 20 μM ZnCl_2_ (for Zn^II^_2_ AdcR) added to these reactions. Anisotropy experiments were performed on an ISS PC1 spectrofluorometer in steady-state mode with Glan-Thompson polarizers in the L-format. The excitation wavelength was set at 494 nm with a 1 mm slit and the total emission intensity collected through a 515 nm filter. For Zn(II)-bound-AdcR DNA-binding experiments, the data were fit with DynaFit (Kuzmic, 1996) using a non-dissociable dimer 1:1 dimer:DNA binding model (*K_dim_*= 10^12^ M^−1^). For Zn(II)-bound experiments, the initial anisotropy (*r*_0_) was fixed to the measured value for the free DNA, with the anisotropy response of the saturated protein:DNA complex (*r_complex_*) optimized during a nonlinear least squares fit using DynaFit (Kuzmic, 1996). Apo binding data were fit in the same manner, except *r_complex_* was fixed to reflect the anisotropy change (*r_complex_* – *r_0_*) observed for wild-type AdcR in the presence of zinc. The errors on *K*_apo,DNA_ and *K*_Zn,DNA_, reflect the standard deviation of 3 independent titrations (Table S2). The coupling free energies were calculated using the following equation: Δ*G_c_*= –*RT*1n(*K*_Zn,DNA_ /*K*_apo,DNA_) (Giedroc & Arunkumar, 2007). Negative values of Δ*G_c_* were observed since AdcR is a positive allosteric activator in the presence of Zn^II^ (*K*_apo,DNA_<*K*_Zn,DNA_,).

## Acknowledgements

We gratefully acknowledge support of this work by the NIH (R35 GM118157 to D. P. G.). NMR instrumentation in the METACyt Biomolecular NMR Laboratory at Indiana University was generously supported by a grant from the Lilly Endowment. D.A.C. acknowledges support from the Pew Latin American Fellows Program in the Biomedical Sciences. We also thank Dr. Lixin Fan of the Small-Angle X-ray Scattering Core Facility, National Cancer Institute, Frederick, MD for acquiring the SAXS data.

## Supplementary Data

**Table S1.**
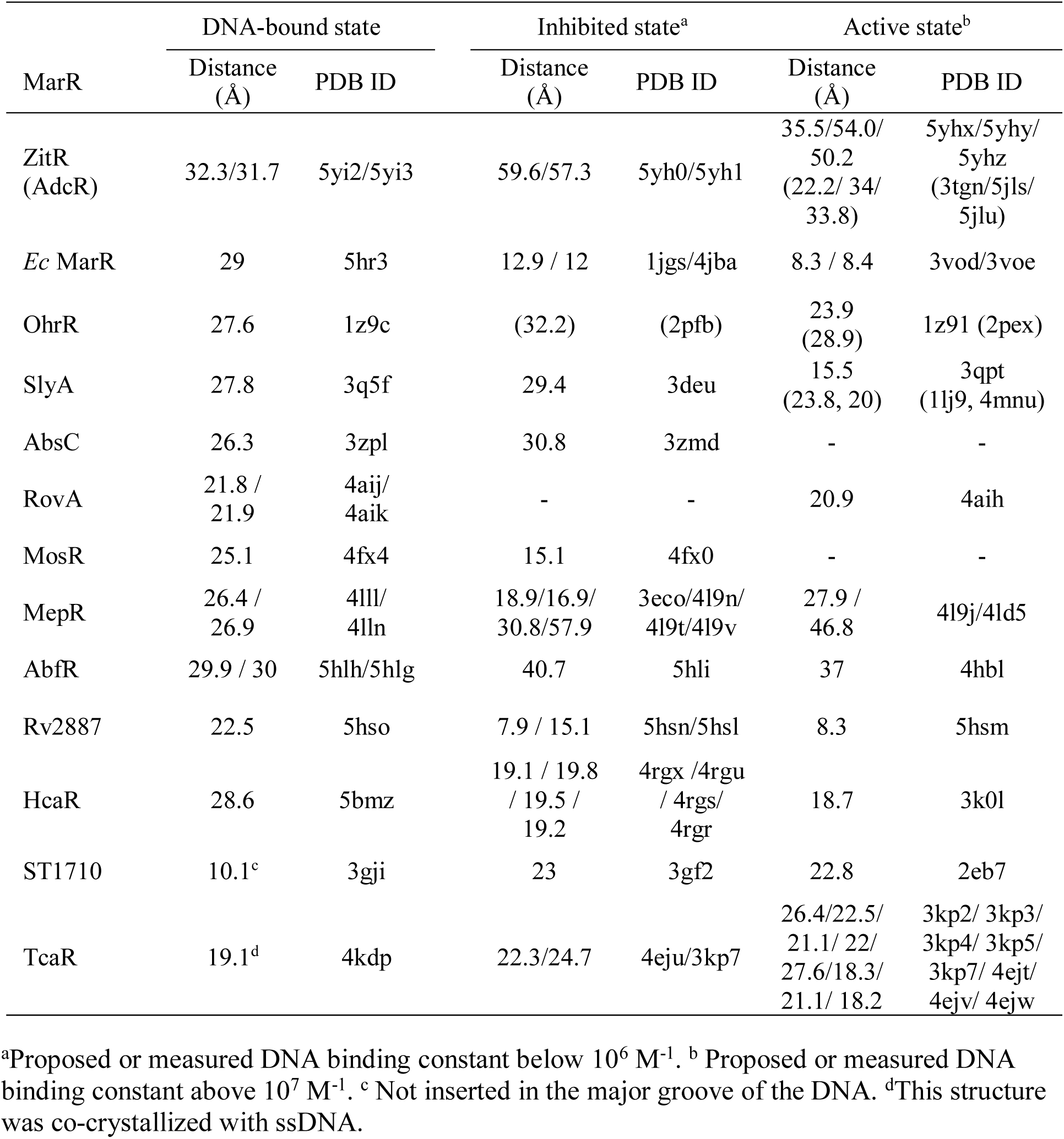
Interprotomer distances between the Cα of the N-terminal residue in the α4 and α4’ helices

| MarR | DNA-bound state |  | Inhibited state<sup>a</sup> |  | Active state<sup>b</sup> |  |
| --- | --- | --- | --- | --- | --- | --- |
|  | Distance (Å) | PDB ID | Distance (Å) | PDB ID | Distance (Å) | PDB ID |
| ZitR (AdcR) | 32.3/31.7 | 5yi2/5yi3 | 59.6/57.3 | 5yh0/5yh1 | 35.5/54.0/<br>50.2<br>(22.2/ 34/<br>33.8) | 5yhx/5yhy/<br>5yhz<br>(3tgn/5jls/<br>5jlu) |
| *Ec* MarR | 29 | 5hr3 | 12.9 / 12 | 1jgs/4jba | 8.3 / 8.4 | 3vod/3voe |
| OhrR | 27.6 | 1z9c | (32.2) | (2pfb) | 23.9<br>(28.9) | 1z91 (2pex) |
| SlyA | 27.8 | 3q5f | 29.4 | 3deu | 15.5<br>(23.8, 20) | 3qpt<br>(1lj9, 4mnu) |
| AbsC | 26.3 | 3zpl | 30.8 | 3zmd | - | - |
| RovA | 21.8 /<br>21.9 | 4aij/<br>4aik | - | - | 20.9 | 4aih |
| MosR | 25.1 | 4fx4 | 15.1 | 4fx0 | - | - |
| MepR | 26.4 /<br>26.9 | 4lll/<br>4lln | 18.9/16.9/<br>30.8/57.9 | 3eco/4l9n/<br>4l9t/4l9v | 27.9 /<br>46.8 | 4l9j/4ld5 |
| AbfR | 29.9 / 30 | 5hlh/5hlg | 40.7 | 5hli | 37 | 4hbl |
| Rv2887 | 22.5 | 5hso | 7.9 / 15.1 | 5hsn/5hsl | 8.3 | 5hsm |
| HcaR | 28.6 | 5bmz | 19.1 / 19.8<br>/ 19.5 /<br>19.2 | 4rgx /4rgu<br>/ 4rgs/<br>4rgr | 18.7 | 3k0l |
| ST1710 | 10.1<sup>c</sup> | 3gji | 23 | 3gf2 | 22.8 | 2eb7 |
| TcaR | 19.1<sup>d</sup> | 4kdp | 22.3/24.7 | 4eju/3kp7 | 26.4/22.5/<br>21.1/ 22/<br>27.6/18.3/<br>21.1/ 18.2 | 3kp2/ 3kp3/<br>3kp4/ 3kp5/<br>3kp7/ 4ejt/<br>4ejv/ 4ejw |

<sup>a</sup>Proposed or measured DNA binding constant below 10<sup>6</sup> M<sup>-1</sup>. <sup>b</sup> Proposed or measured DNA binding constant above 10<sup>7</sup> M<sup>-1</sup>. <sup>c</sup> Not inserted in the major groove of the DNA. <sup>d</sup>This structure was co-crystallized with ssDNA.

**Table S2.** DNA binding parameters for wild-type AdcR and substitution mutants^a^

| AdcR | $K_{apo,DNA}$<br>($\times 10^6 M^{-1}$) | Zn<sup>II</sup> | | Dynamic changes (Zn<sup>II</sup>) | | Fractional<br>ASA<sup>b</sup> |
| --- | --- | --- | --- | --- | --- | --- |
| | | $K_{Zn,DNA}$<br>($\times 10^6 M^{-1}$) | $\Delta G_c$<br>(kcal mol<sup>-1</sup>) | $\Delta S^2_{axis}$ | $\Delta R_{ex}$ | |
| wild-type | 0.5 ± 0.2 | 450 ± 220 | -4.0 ± 0.6 |  |  |  |
| I104A | 0.20 ± 0.01 | 280 ± 30 | -4.3 ± 0.4 | -0.08 ± 0.01 | 0.3 | 0.04 |
| L36A | 0.07 ± 0.01 | 80 ± 30 | -4.1 ± 0.4 | 0.15 ± 0.02 | 3.2 | 0.05 |
| V34A | 0.5<sup>c</sup> ± 0.01 | 13 ± 1 | -2.2 ± 0.3 | 0.15 ± 0.02 | 3.5 | 0.46 |
| L81V | 0.16 ± 0.12 | 12 ± 8 | -2.4 ± 0.6 | 0.20 ± 0.02 | 0 | 0.00 |
| L61V | 0.44 ± 0.05 | 21 ± 18 | -2.3 ± 0.1 | -0.247 ±<br>0.001 | -0.7 | 0.01 |
| L57M | 0.035 ±<br>0.030 | 1 ± 0.2 | -2.0 ± 0.7 | -0.30 ± 0.02 | 0.7 | 0.00 |
| L57V | <0.05<sup>d</sup> | <0.05<sup>d</sup> | N/A | -0.30 ± 0.02 | 0.7 | 0.00 |
| I16A | 1.8 ± 0.9 | 17 ± 14 | -1.8 ± 0.4 | -0.07 ± 0.01 | 3.5 | 0.11 |
| L4A | 0.5 ± 0.2 | 11 ± 8 | -1.8 ± 0.3 | 0.02 ± 0.02 | 4 | 0.01 |
| V142A | 0.41 ± 0.05 | 4.1 ± 2.3 | -1.4 ± 0.2 | -0.095 ±<br>0.005 | 7 | 0.31 |
| N38A | 0.05 ± 0.01 | 19 ± 10 | -3.5 ± 0.7 | —<sup>e</sup> | — | — |
| N38A/Q40A | 0.10 ± 0.04 | 2.2 ± 0.4 | -1.9 ± 0.2 | — | — | — |
| E24D | 0.17 ± 0.04 | 2.2 ± 1.7 | -1.6 ± 0.3 | — | — | — |

<sup>a</sup>Conditions: 10 mM Hepes, pH 7.0, 0.23 M NaCl, 1 mM TCEP (chelexed), 10 nM DNA, 25.0 °C with 2.0 mM EDTA (for apo-AdcR) or 20 μM ZnCl<sub>2</sub> (for Zn<sup>II</sup><sub>2</sub> AdcR) added to these reactions. See Fig. 6C, main text, for a graphical representation of these data. <sup>b</sup>Accessible surface area (ASA) calculated from the Zn<sup>II</sup><sub>2</sub>-bound AdcR (Guerra *et al.*, 2011). <sup>c</sup>Upper limit on measurable $K_{apo,DNA}$ under these solution conditions. <sup>d</sup>Weaker than upper limit. <sup>e</sup>Not measurable using the NMR experiments employed here.

**Table S3.**
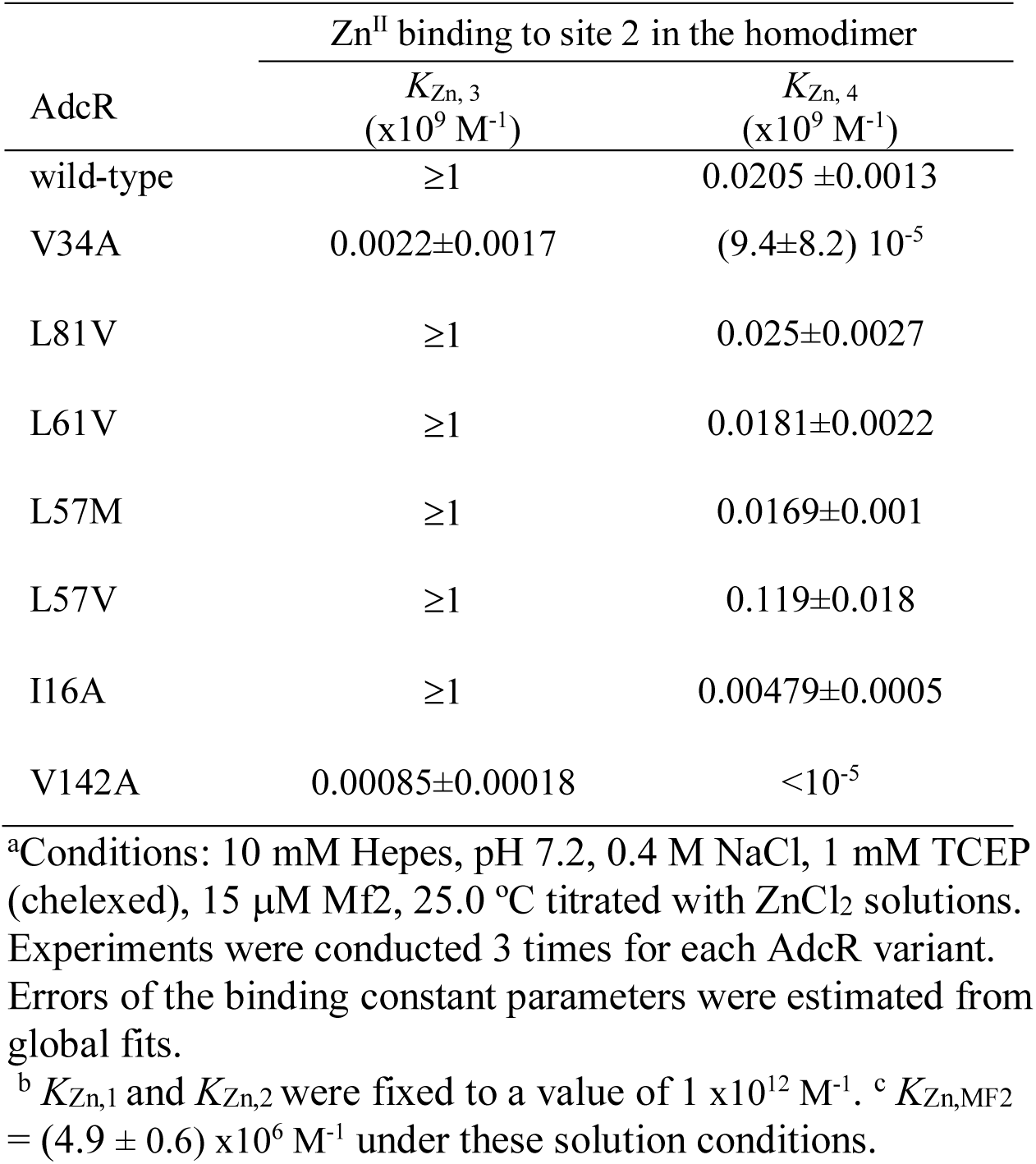
Zinc binding affinities of wild-type AdcR and selected AdcR mutants characterized here.

**Figure S1.**
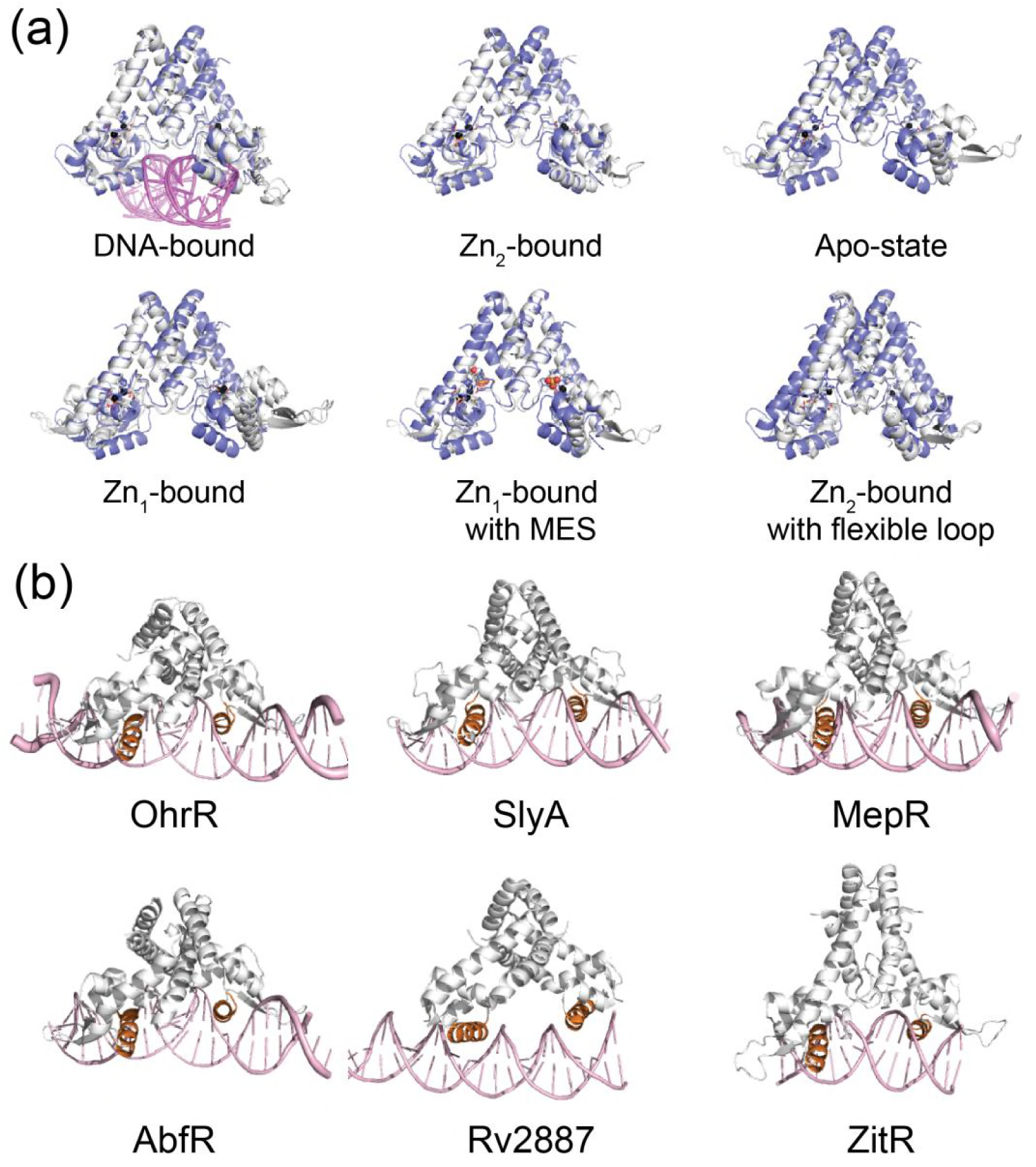
Structural comparison (global superposition) between AdcR (3tgn, shaded *slate* in all panels, (Guerra *et al.*, 2011)) and *L. lactis* ZitR (Zhu *et al.*, 2017) in the different allosteric states (DNA-bound PDB codes, 5yi2, 5yi3; Zn^II^_2_-bound, 5hyx; Apo-state, 5yi1; Zn_1_-bound PDB ID 5yhy, 5yl0; Zn^II^_2_-bound alternative state with a MES molecule in Zn site 1, 5yhz; Zn_2_-bound from Group A *Streptococcus pyogenes* AdcR (Sanson *et al.*, 2015) with flexible loop, 5jls, 5lju). (b) Structural comparisons of various MarR family repressors in the DNA-bound states. (*B. subtilis* OhrR, PDB code, 1z9c; *S. enterica* SlyA, 3q5f; *S. aureus* MepR, 4lln; *S. epidermis* AbfR, 5hlg; *M. tuberculosis* Rv2887, 5hso; *L. Lactis* ZitR, 5yi2.

**Figure S2.**
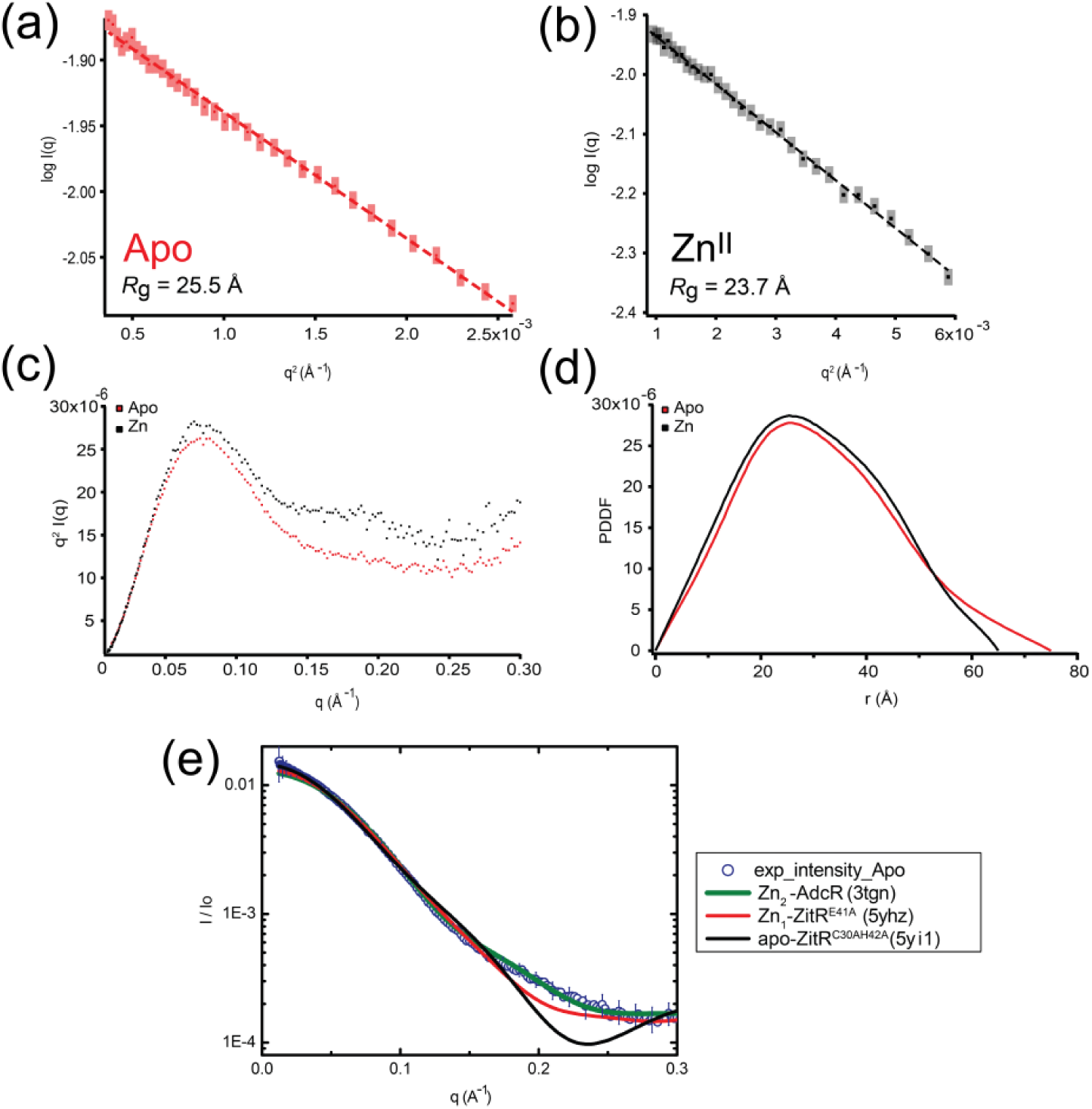
Small angle X-ray scattering (SAXS) analysis of AdcR in apo and Zn-binding states. (a) The Guinier region with linear fit of the scattering curve of the AdcR apo state (red) and Zn-binding state (black). Radius of gyration (Rg) of each state is presented at the low left corner. Note that scattering intensity is in arbitrary unit. (b) Dimensolionless Kratky plot of the AdcR apo (red) and Zn-binding (red) state. (c) Pair distance distribution function (PDDF) of the AdcR apo (red) and Zn-binding (black) state. The end-to-end distance (*D*_max_) of apo state is 65 Å and *D*_max_ of the Zn-binding state is 75 Å. (d) Scattering profiles of AdcR apo (red) and Zn-binding states and (e) calculated scattering profiles of crystal structures of apo ZitR^C30AH42A^ (5yi1) and Zn-binding ZitR^E41A^ (5yhz) (Zhu *et al.*, 2017) compared to the experimental and fitted curves obtained for apo-AdcR (see **Fig. 2**, main text).

**Figure S3.**
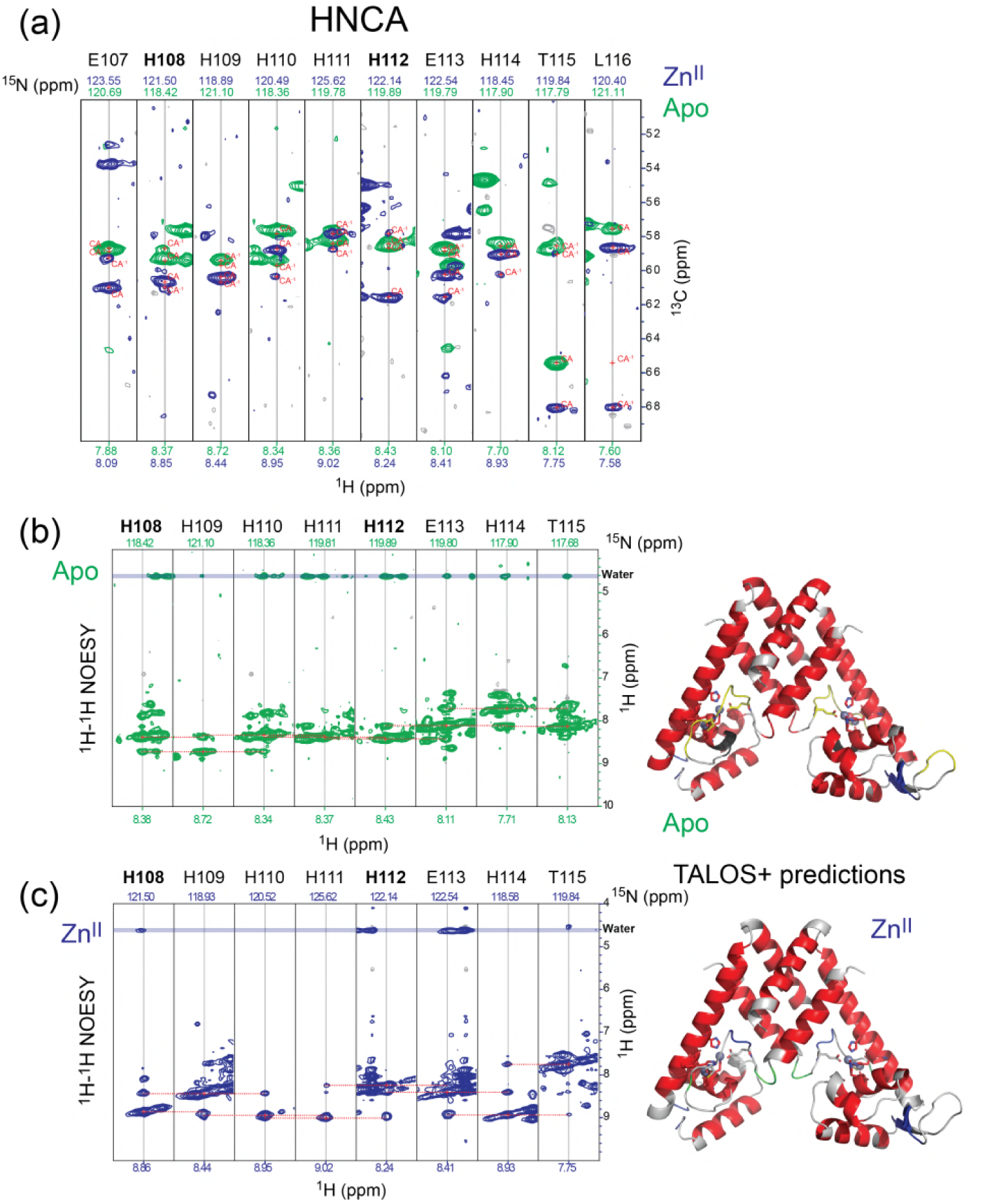
(a) Sequential residue-specific connectivities that link the chemical shifts of the ^13^Cα resonances in the α5 helix (E107-L116; H108, H112 Zn^II^ ligands in bold) from an HNCA experiment. (b), (c) ^1^H,^15^N NOESY-HSQC strips obtained from the same region of the α5 helix in the apo- (b) and Zn^II^_2_ (c) states. *Right*, TALOS+ predictions in the apo- (top) and Zn^II^_2_ (bottom) states. Despite fully α-helical predictions, the apo-state is characterized by weaker i, *i*+*2* NH-NH correlations, and stronger NOEs (as solvent exchange crosspeaks) to water, relative to the Zn^II^ state. This is consistent with a more highly dynamic α5 helix in the apo-state.

**Figure S4.**
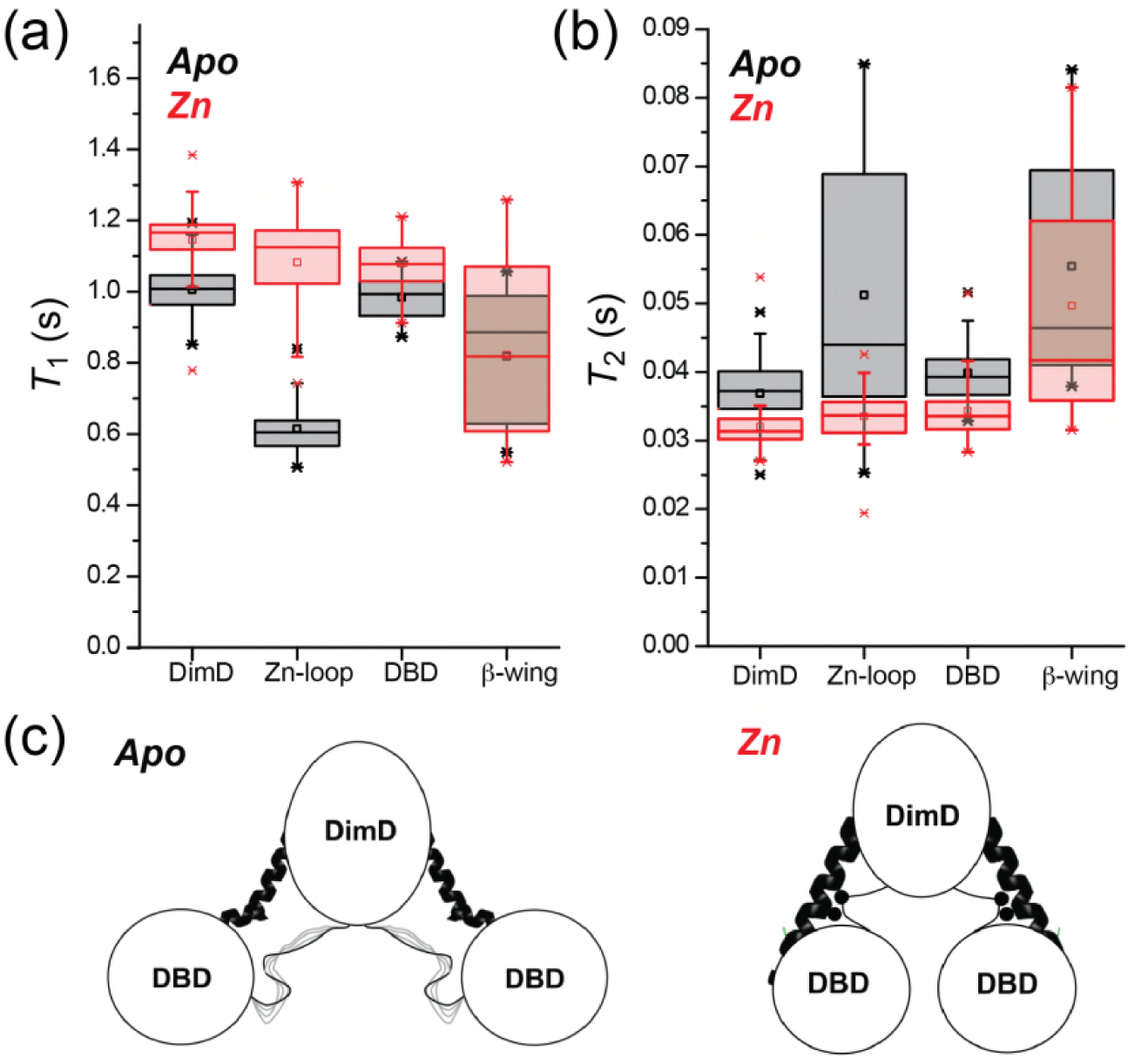
Average backbone amide ^1^H-^15^N relaxation parameters *T*_1_ (1/*R*_1_, a) and *T*_2_ (1/*R*_2_, b) for the apo- (*black* boxes) and Zn^II^_2_- (*red* boxes) states of the AdcR homodimer in different regions of the molecule: DimD, dimerization domain (residues 5-20, 101-144); Zn-loop (α1-α2 loop, residues 21-37); DBD, DNA binding domain (residues 38-101, excluding the β-wing; β-wing (residues 81-101). (c) Cartoon representations of the data shown in panels (a) and (b) in which the two linkers that connect the two domains (middle of the α5-helix, *dark coil*; α1-α2 loop, *light pencil*) are more dynamic in the apo-state. Note that residues analogous to the α1-α2 loop in AdcR are not observed in the crystal structure of the apo-state of *L. lactis* ZitR (Zhu *et al.*, 2017) (see Fig. S1A), consistent with these findings in solution in apo-AdcR.

**Figure S5.**
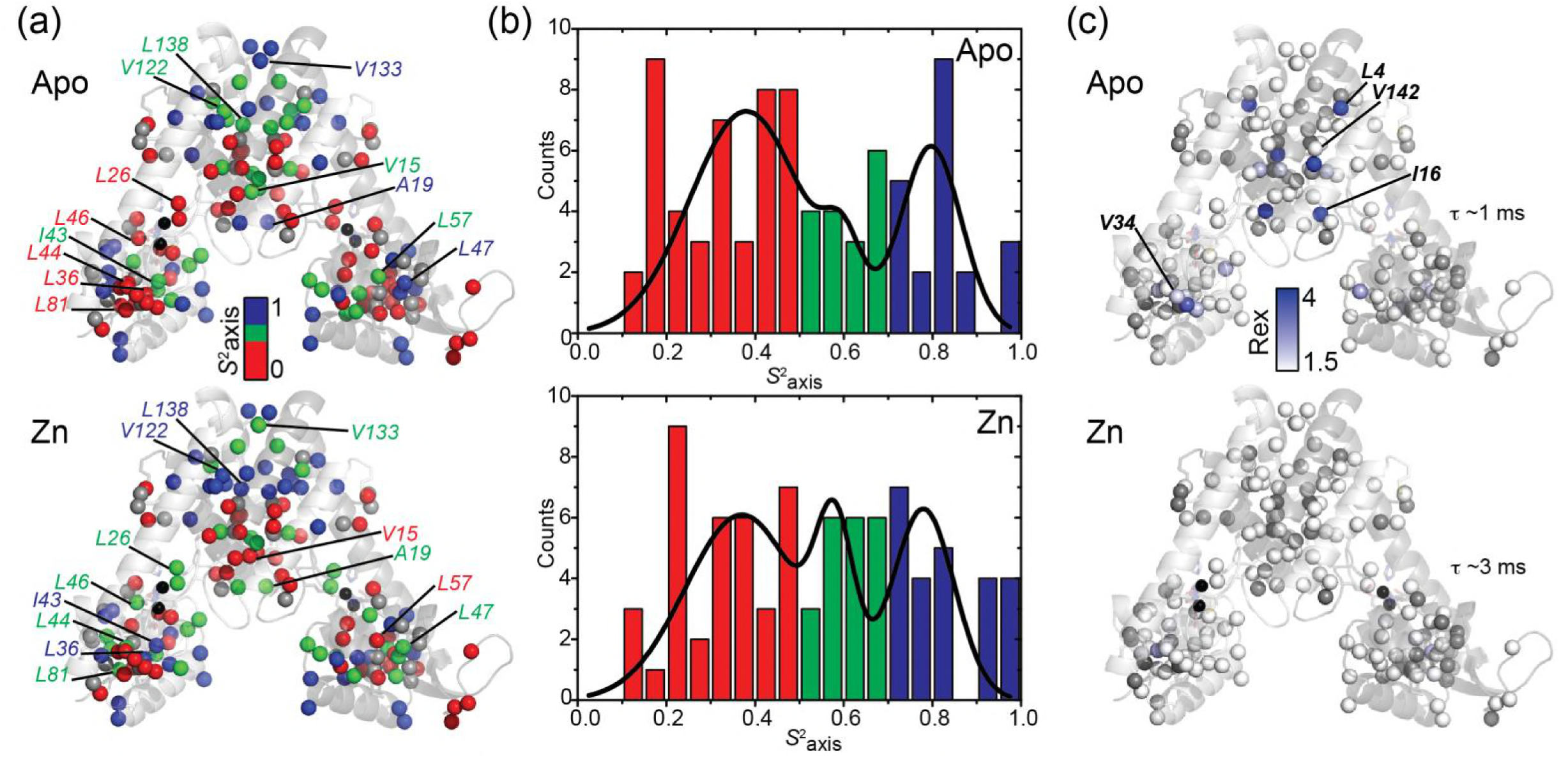
Absolute values of methyl group order parameters, *S*^2^_axis_ (a) and *R*_ex_ (c) on the methyl-bearing residues. (b) Histogram plot of *S*^2^_axis_ from fitting the apo (*top*) and Zn^II^_2_ (*bottom*) states in panel (a) calculated according to (Marlow *et al.*, 2010).

**Figure S6.**
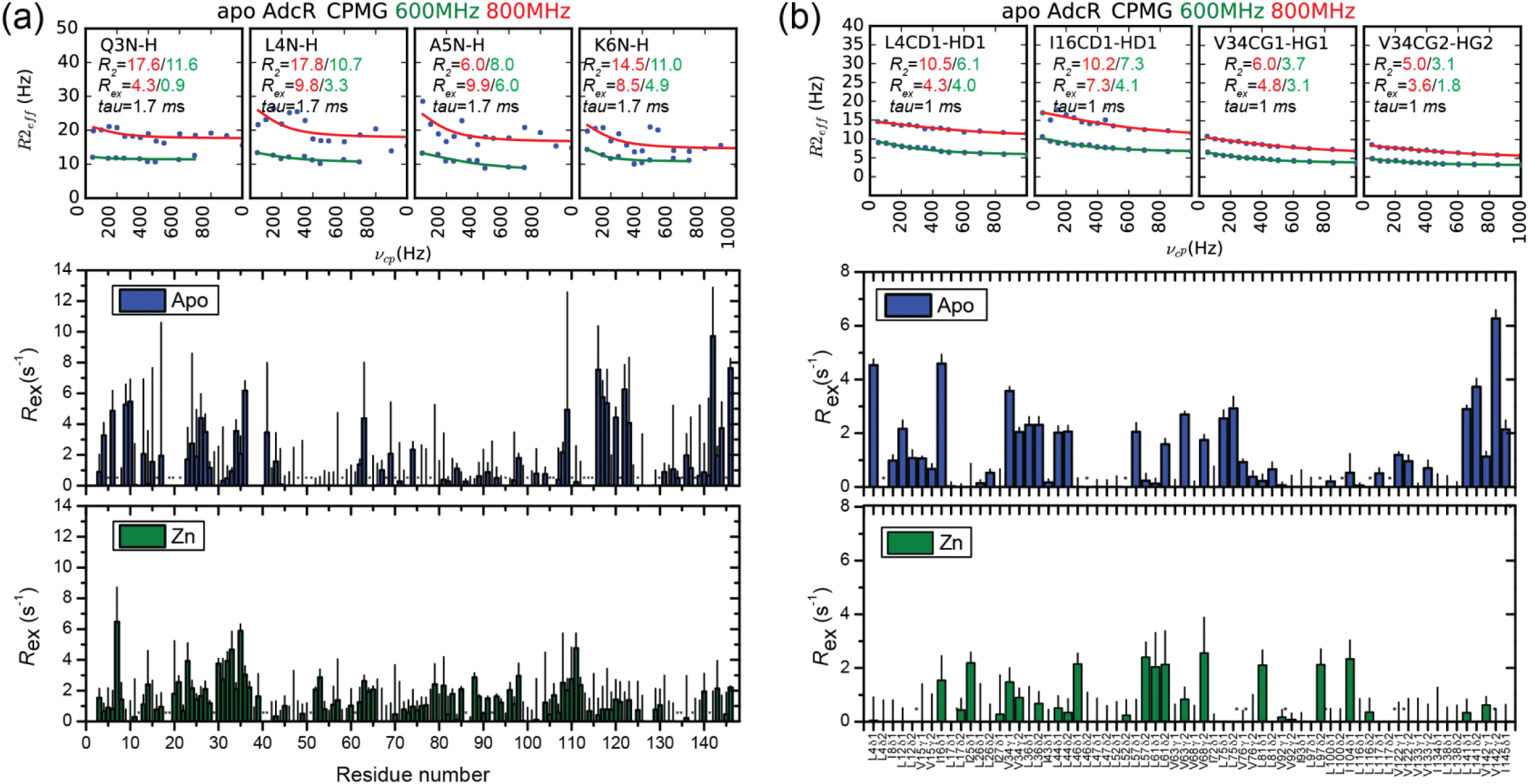
Representative raw relaxation dispersion curves obtained for the indicated backbone (NH) (a) or methyl group (b) used to obtain *R*_ex_ at 600 MHz and 800 MHz. *R*_ex_ for both allosteric states at 600 MHz are shown in each panel. All the residues excluded due to overlap are shown with an asterisk (Backbone Apo: 5, 7, 16, 18, 19, 22, 48, 50, 51, 58, 64, 68, 70, 73, 75, 78, 95, 100, 105, 106, 110, 113, 114, 115, 121, 125, 130, 135, 138, 145; Backbone Zn-bound: 12, 18, 19, 29, 40, 41, 51, 61, 86, 92, 134, 135, 137, 141; Sidechain Apo: L4-δ2, L46-δ2, L52-δ2, L97-δ2, L100-δ2, L116-δ2, L117-δ2; Sidechain Zn-bound: L12-δ2, L17-δ1,L57-δ1, L75-δ1, L75-δ2, L81-δ2, L97-δ1, L117-δ1, L117-δ2, L141-δ2).

**Figure S7.**
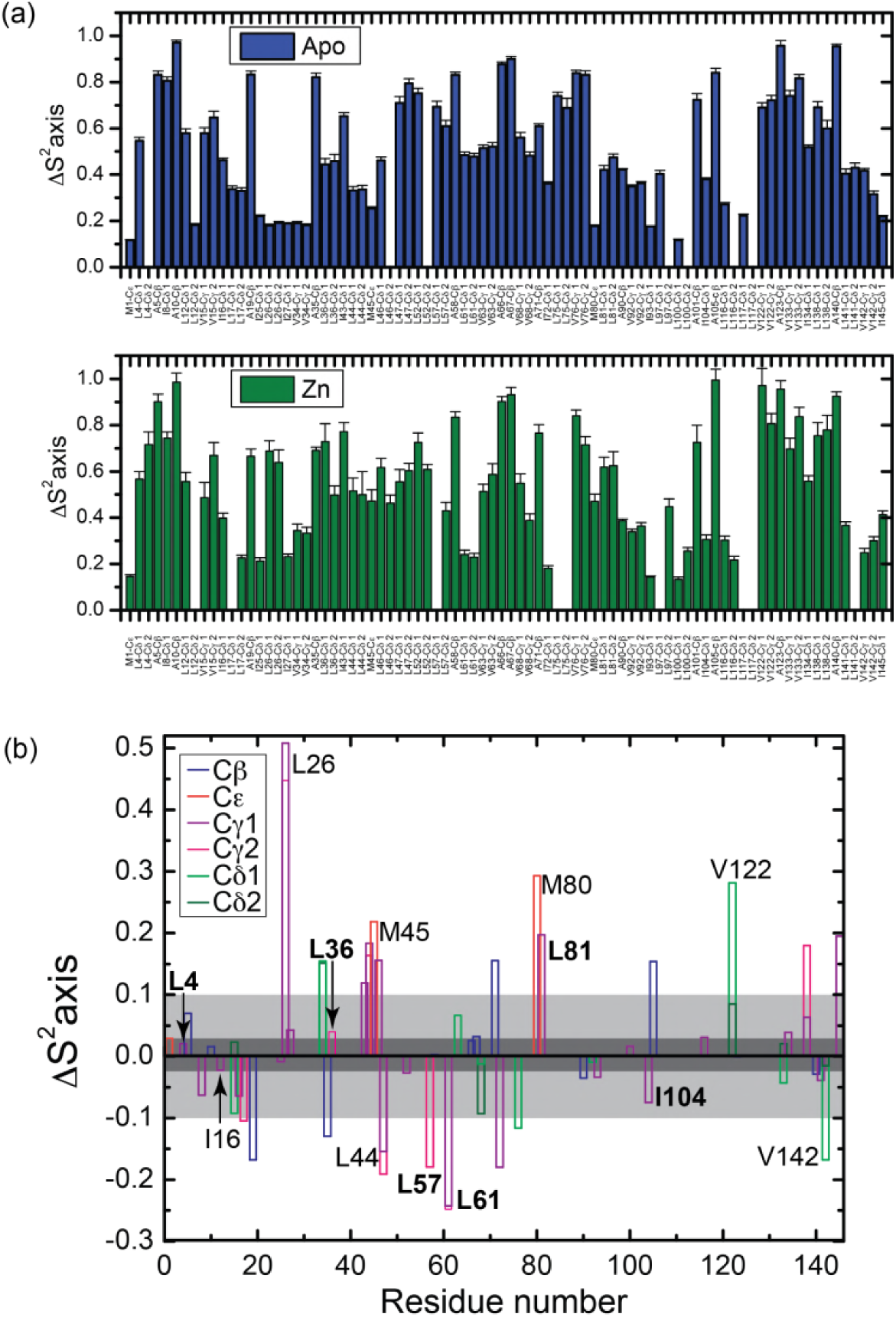
(a) Stereospecific methyl group axial order parameters, *S*^2^_axis_, for apo- and Zn^II^_2_- AdcR as measured at 600 MHz (similar results were obtained at 800 MHz; data not shown). (b) Difference in axial order parameter (Δ*S*^2^_axis_= *S*^2^_axis_^Zn^–*S*^2^_axis_^apo^) between apo- and Zn^II^_2_-states, with the specific type of methyl group color-coded as indicated: Cβ, Ala; Cε, Met; Cγ1, Cγ2, Val; Cδ1, Cδ2, Leu.

**Figure S8.**
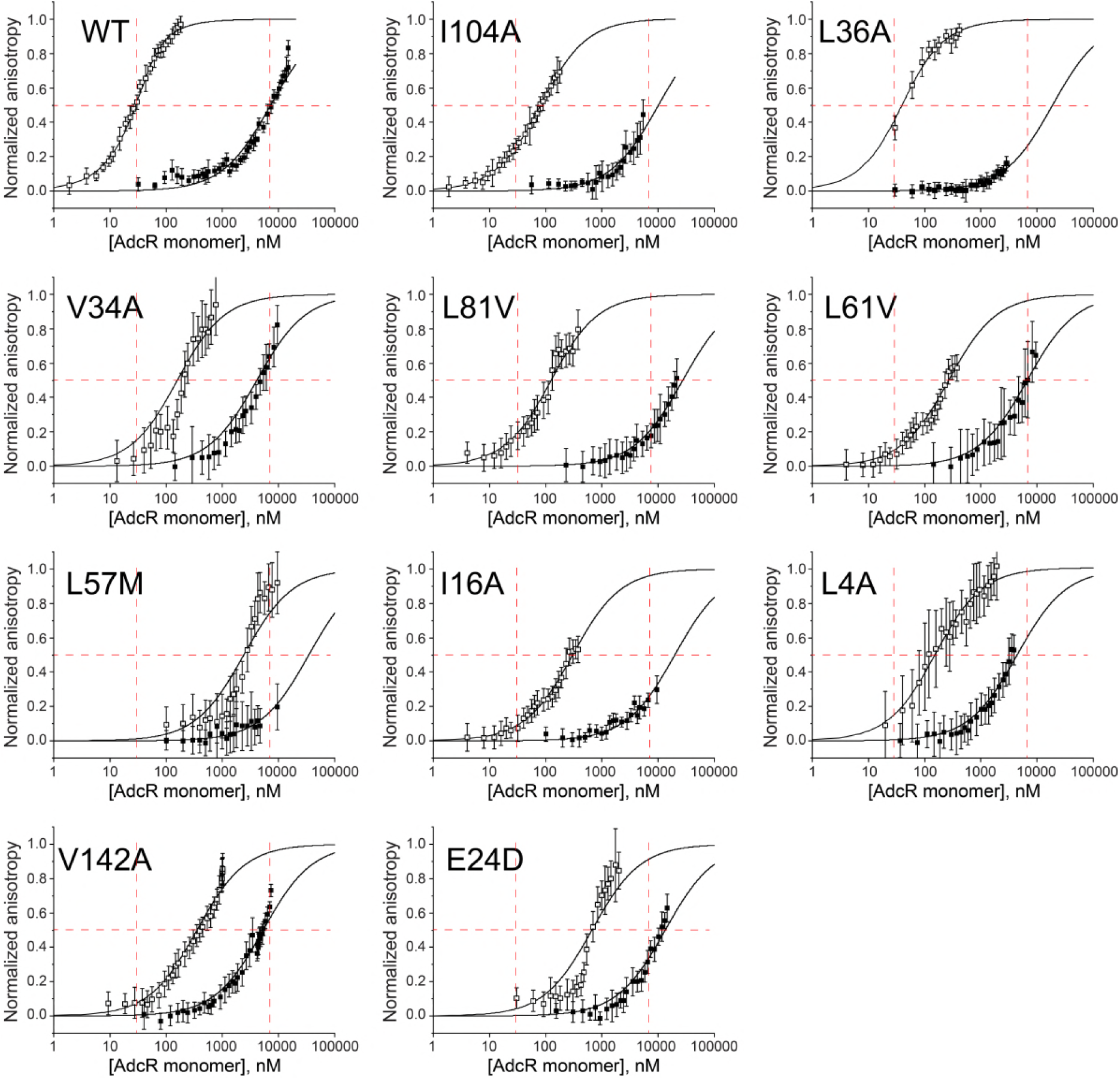
Representative DNA operator binding isotherms obtained for selected for wild-type (WT) AdcR and selected AdcR mutants in the apo- and Zn^II^_2_-states. The *continuous* lines through each set of data correspond to nonlinear least squares fit to a 1:1 non-dissociable AdcR dimer binding model, with parameters compiled in Table S2, and Δ*G*_c_ shown graphically in Fig. 6c (**main text**). The *red vertical and horizontal lines* represent the AdcR monomer concentrations that correspond to 50% DNA-saturation points for the wild-type AdcR under the same solution conditions, presented as a guide only. Conditions: 10 mM Hepes, pH 7.0, 0.23 M NaCl, 1 mM TCEP (chelexed), 10 or 20 nM nM DNA, 25.0 °C with 1.0 mM EDTA (for apo- AdcR) or 20 μM ZnCl_2_ (for Zn^II^_2_ AdcR) added to these reactions. Experiments were conducted 3 times for each AdcR variant.

**Figure S9.**
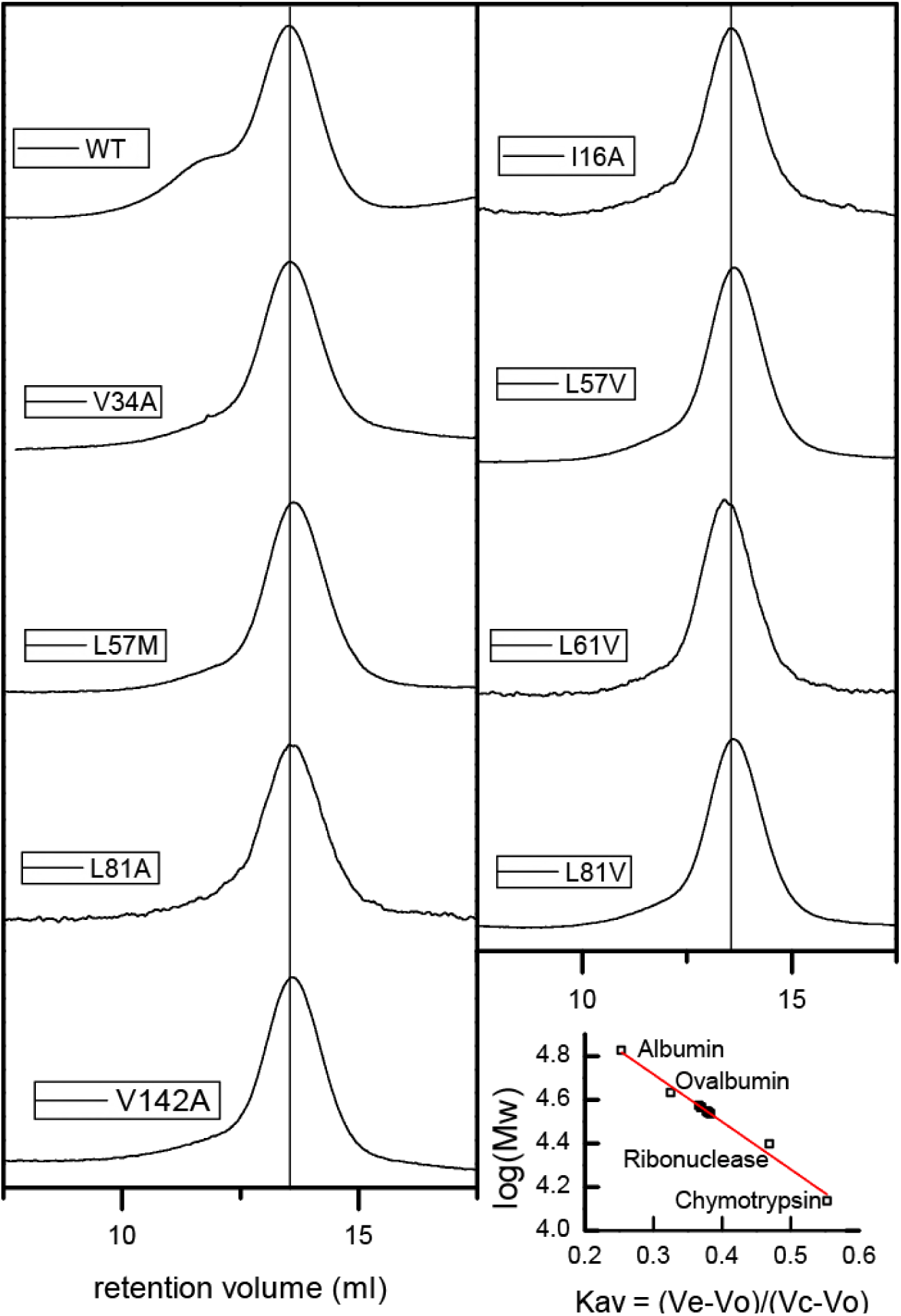
Gel filtration chromatograms for AdcR variants in the apo-state. *Lower right*, calibration curve with standards (empty squares) and AdcR variants (*filled* squares).

**Figure S10.**
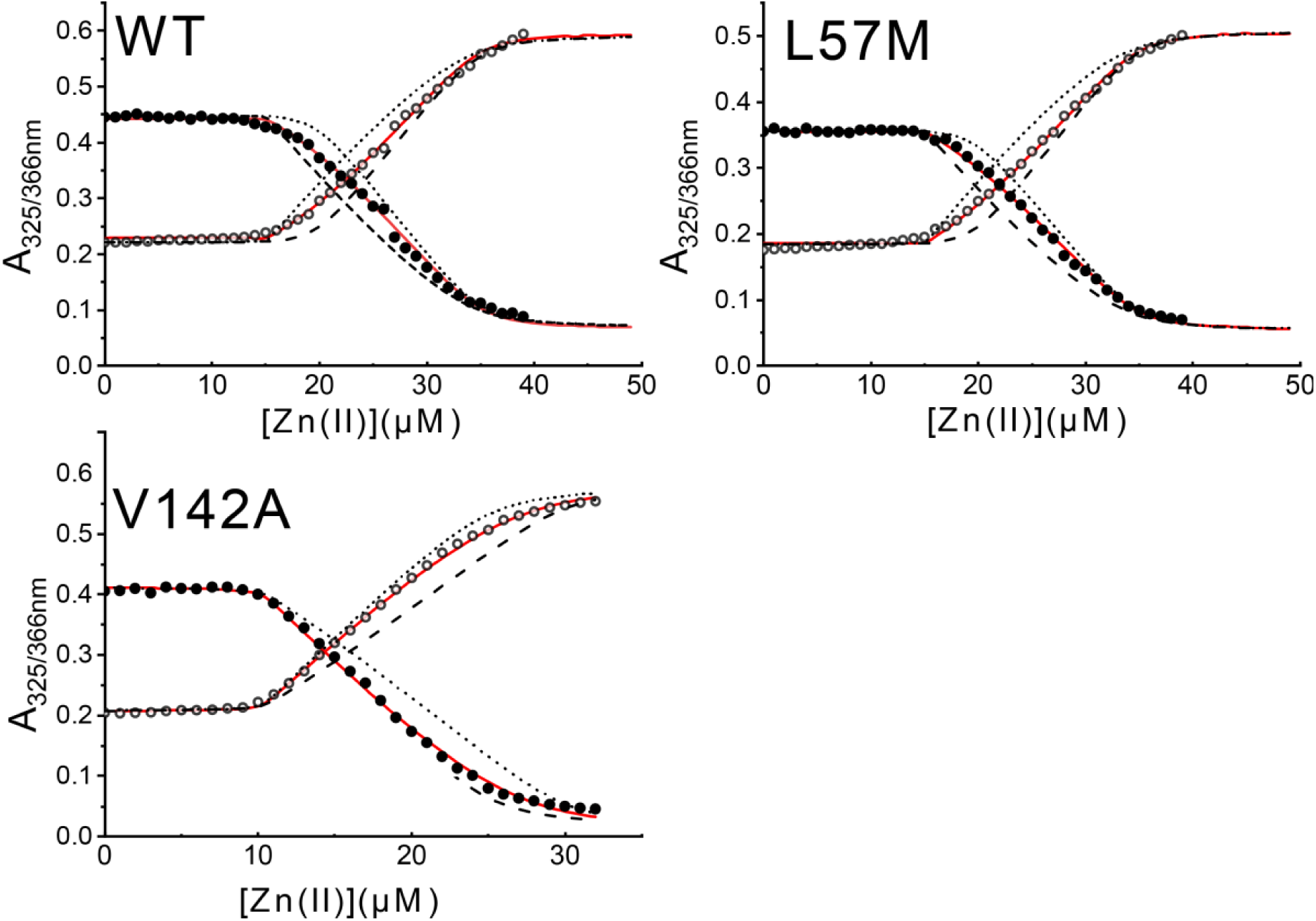
Representative Zn^II^-binding isotherms obtained from a titration of apo (metal-free) wild-type AdcR or a mutant AdcR and mag-fura-2 (mf2) with ZnSO_4_. Zn^II^ binding parameters for these and other AdcRs are compiled in Table S3. Experiments were conducted 3 times for each AdcR variant.

**Figure S11.**
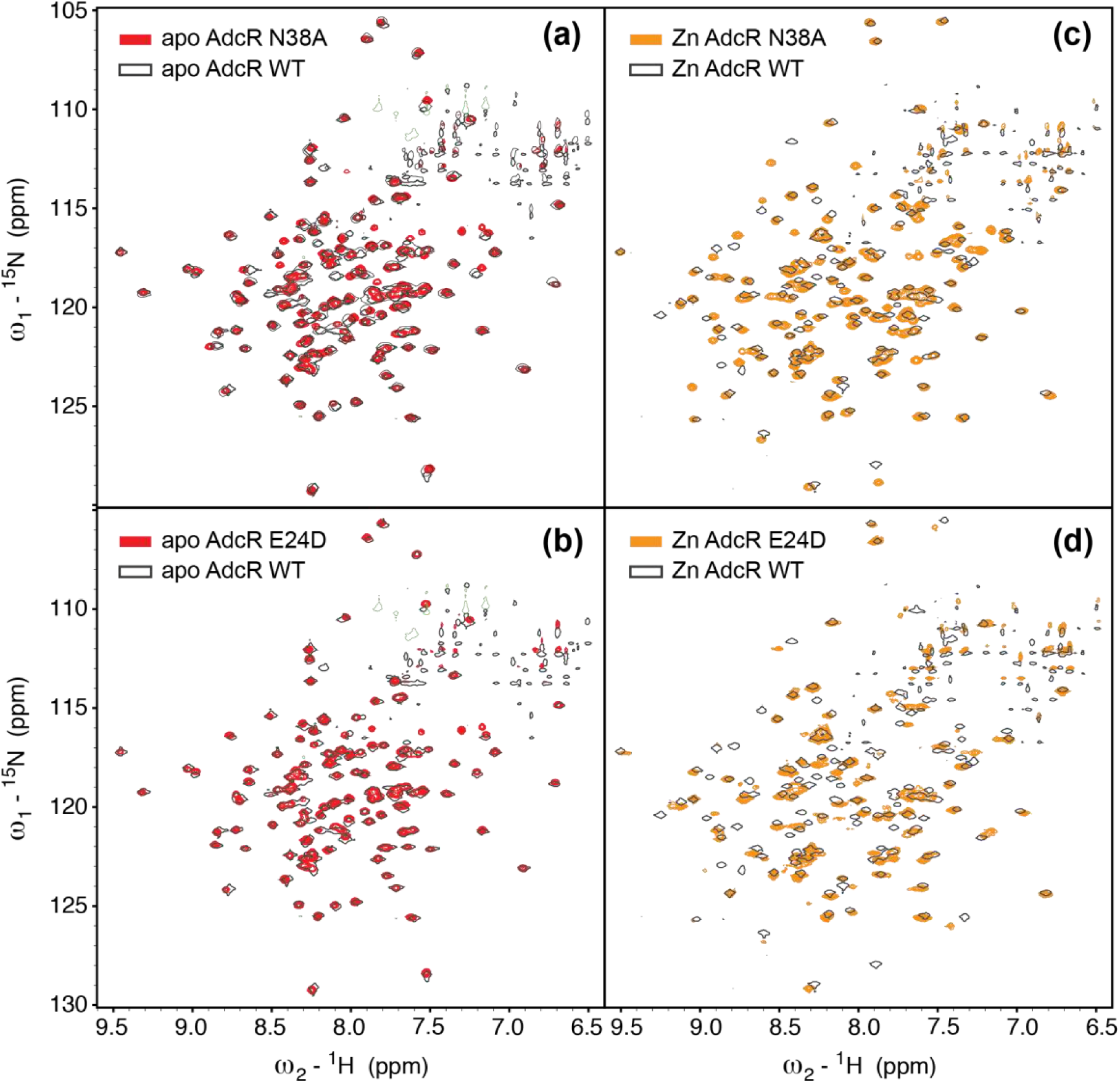
2D ^1^H,^15^N TROSY spectra of apo- (*left*) and Zn^II^_2_ (*right*) states of N38A and E24D AdcRs, compared to the wild-type AdcR (*black* contour; since contour line shown) acquired under the same solution conditions (50 mM NaCl, pH 6.0, 35 °C).

**Figure S12.**
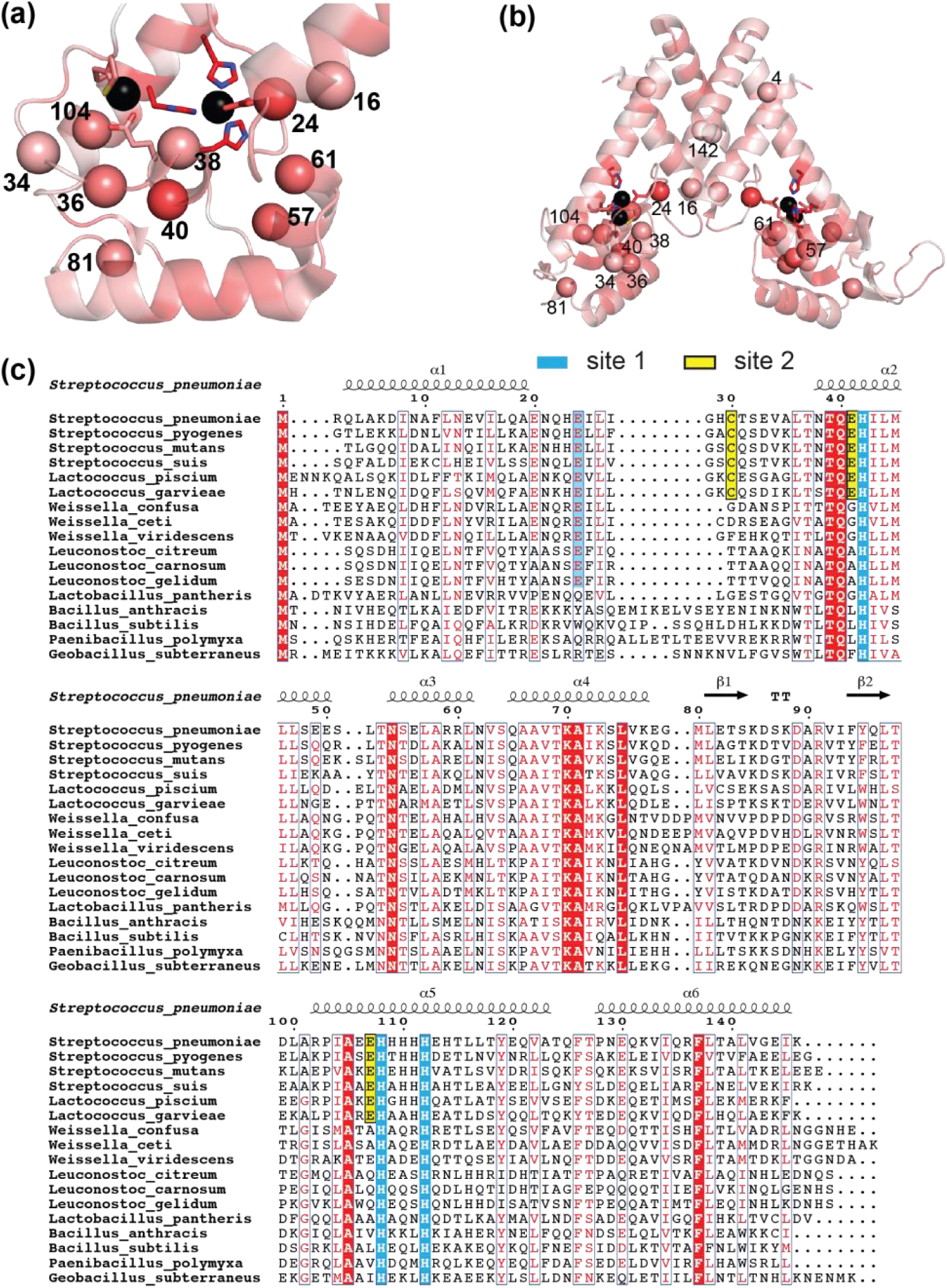
Amino acid sequence conservation of *S. pneumoniae* AdcR and candidate closely related MarR family repressors. Sequence conservation highlighting those residues targeted for mutagenesis in this work with a Cα sphere on the Zn^II^_2_ AdcR structure (Guerra *et al.*, 2011) in the DNA binding domain (a) and the entire molecule (b). The ribbon structure shows the degree of conservation by ramping the color from *white* to *bright red*, with those residues of high conservation shaded *bright red*, using Protskin (Ritter *et al.*, 2004). For reference, Zn^II^ ligands are invariant (100% conserved). (b) Multiple sequence analysis of the 17 AdcR-like repressors used to create the sequence conservation map.

